# Mitochondrial COA7 is a heme-binding protein involved in the early stages of complex IV assembly

**DOI:** 10.1101/2021.06.10.447992

**Authors:** Luke E. Formosa, Shadi Maghool, Alice J. Sharpe, Boris Reljic, Linden Muellner-Wong, David A. Stroud, Michael T. Ryan, Megan J. Maher

**Author notes:** These authors contributed equally to this work (Luke E. Formosa, Shadi Maghool). **Author Contributions** LEF, SM, MJM and MTR designed research; LEF, SM, AJS, BR, LM-W, DAS, MJM and MTR performed research; LEF, SM, BR, DAS, MJM and MTR analyzed data; LEF, SM, MJM and MTR wrote the paper. **Competing Interests** Statement The authors declare no competing interests.

## Abstract

Cytochrome *c* oxidase assembly factor 7 (COA7) is a metazoan-specific assembly factor, critical for the biogenesis of mitochondrial complex IV (cytochrome *c* oxidase). Although mutations in COA7 have been linked in patients to complex IV assembly defects and neurological conditions such as peripheral neuropathy, ataxia and leukoencephalopathy, the precise role COA7 plays in the biogenesis of complex IV is not known. Here we show that the absence of COA7 leads to arrest of the complex IV assembly pathway at the initial step where the COX1 module is built, which requires incorporation of copper and heme cofactors. In solution, purified COA7 binds heme with micromolar affinity, through axial ligation to the central iron atom by histidine and methionine residues. Surprisingly, the crystal structure of COA7, determined to 2.4 Å resolution, reveals a ‘banana-shaped’ molecule composed of five helix-turn-helix (α/α) repeats, tethered by disulfide bonds, with a structure entirely distinct from proteins with characterized heme binding activities. We therefore propose a role for COA7 in heme binding/chaperoning in the mitochondrial intermembrane space, this activity being crucial for and providing a missing link in complex IV biogenesis.

**Significance Statement:** Assembly factors play key roles in the biogenesis of many mitochondrial protein complexes regulating their stability, activity and incorporation of essential cofactors. COA7 is a metazoan-specific assembly factor, the absence or mutation of which in humans accompanies complex IV assembly defects and neurological conditions. Here we report the crystal structure of COA7 to 2.4 Å resolution, revealing a ‘banana-shaped’ molecule composed of five helix-turn-helix (α/α) repeats, tethered by disulfide bonds. Characterization of pathogenic variants reveals significantly lower stabilities, correlating with the associated disease outcomes. Fascinatingly, COA7 binds heme with micromolar affinity, despite the fact that the protein structure does not resemble previously characterized heme-binding proteins. This provides a possible missing link for heme handling in the mitochondrial intermembrane space.

## Introduction

In eukaryotes, the bulk of ATP generation occurs through oxidative phosphorylation (OXPHOS) that involves five multi-subunit protein complexes in the inner mitochondrial membrane (IMM), known as complexes I to V (1). Complex IV (cytochrome *c* oxidase or COX) is the copper and heme *a* containing terminal oxidase of the mitochondrial respiratory chain, which catalyzes electron transfer from cytochrome *c* in the intermembrane space (IMS) to molecular oxygen and contributes to the generation of the proton gradient required to power ATP synthesis. In mammals, complex IV is composed of 14 subunits, including three core subunits (COX1, COX2 and COX3) that are encoded by mitochondrial DNA (mtDNA) (2-4).

The assembly of complex IV is complicated, but is thought to involve the biogenesis of different modules harboring both mtDNA and nuclear encoded subunits (5, 6). A central module is seeded by membrane integration and maturation of COX1 that then incorporates separate modules containing COX2 and COX3. The incorporation of three copper ions, heme *a* and heme *a*_3_ cofactors is necessary for the biogenesis of the COX1 and COX2 subunits and assembly of complex IV. In addition, complex IV requires the additional action of more than 30 assembly factors (6). While not part of the final enzyme, these assembly factors have been shown to perform critical roles in subunit maturation, co-factor incorporation and stabilization of intermediate assemblies of complex IV in humans. Loss-of-function mutations in both mtDNA and nuclear DNA genes encoding mitochondrial proteins, including complex IV subunits and assembly factors, lead to complex IV deficiency and mitochondrial disease. Complex IV is also found in a number of supercomplexes, which are large assemblies containing complex IV along with complexes I and III, including the complex I-complex III_2_-complex IV (CI/III_2_/IV) respirasome (7-9).

The cytochrome *c* oxidase assembly factor 7 (COA7, also referred to as RESA1) is an IMS localized (10, 11), metazoan-specific complex IV assembly factor. The function of COA7 in complex IV assembly is unknown. COA7 is a cysteine-rich protein (13 cysteine residues) which based on sequence analysis, contains five Sel1-like repeat domains. Sel1-like repeat (SLR) proteins are a sub-class of the tetratricopeptide repeat (TPR) proteins, which belong to the α/α repeat family of solenoid proteins (12, 13). Available structures of SLR proteins show large solvent accessible surfaces, suitable for the binding of both large and small substrates (12-14). Accordingly, solenoid proteins often function in DNA-, peptide- and protein-protein interactions (12-14), which facilitate cellular processes such as signaling, the cell-cycle and inter-compartmental transport (13, 15-17).

A number of pathogenic mutations in the gene encoding COA7 have been identified in patients exhibiting neurological symptoms of peripheral neuropathy, ataxia and leukoencephalopathy (18, 19). The first of these studies described an isolated deficiency of complex IV in patient skin fibroblasts and skeletal muscle in the presence of compound heterozygous mutations in *COA7* (p. Tyr137Cys and COA7-exon2Δ), resulting in the expression of the COA7 mutant proteins Y137C and an in-frame deletion mutant, missing residues 37-83. A second study examined a cohort of Japanese patients with recessive mutations in *COA7*, where skin fibroblasts from these patients showed significant decreases in complex I and complex IV activities. Patients had either homozygous single amino acid p.Asp6Gly (D6G) mutations, compound D6G with p.Ser149Ile (S149I) or a missense p.Gly144fs mutation, and compound p.Arg39Trp (R39W) and COA7-exon2Δ mutations (19). The Arg39, Tyr137 and Ser149 residues are highly conserved among COA7 proteins from different species (**Fig. S1**).

In the present study, we used structural and functional approaches to investigate the role of COA7 in mitochondrial complex biogenesis. Loss of COA7 blocks complex IV assembly steps subsequent to formation of the COX1 intermediate module, with COX2 being highly labile. We report the crystal structure of human COA7, to 2.4 Å resolution, which allows us to propose the molecular origins of the pathogeneses observed for the patient mutations. Furthermore, characterization of recombinant COA7 shows that the protein binds heme *in vitro* specifically and with micromolar affinity, through axial His/Met coordination to the iron atom, despite no resemblance between the COA7 structure and well characterized heme-binding proteins. These findings point to a role for COA7 in binding and/or incorporation of heme into complex IV.

## Results and Discussion

### Loss of COA7 results in reduced levels of respiratory chain enzymes, with a severe complex IV assembly defect

In order to investigate the function of COA7 in the biogenesis of the respiratory chain, CRISPR-Cas9 mediated gene editing (20) was used to generate COA7^KO^ HEK293T cell lines. Following transfection, cells were sorted to generate individual clones. Two clones (termed COA7^KO^-1 and COA7^KO^-2) were found to contain frameshift deletions in the *COA7* gene where the gRNA was targeted. Mitochondria were isolated from both control and COA7^KO^ cells, solubilized in digitonin and subjected to blue native polyacrylamide gel electrophoresis (BN-PAGE) and immunoblotted for components of the respiratory chain complexes (**Fig. 1*A***). Probing for the complex IV subunit COX4, revealed a severe complex IV defect, with no complex IV holoenzyme nor higher complex IV-containing supercomplexes (CI/III_2_/IV or III_2_/IV) detected (**Fig. 1*A***). Consistent with this, probing for the complex I subunit NDUFA9 revealed the loss of the respirasome (CI/III_2_/IV) while a faster migrating species corresponding to the CI/III_2_ supercomplex was seen (**Fig. 1*A***). Probing for the complex III subunit UQCRC1, revealed the loss of the III_2_/IV supercomplex and the accumulation of the CI/III_2_ supercomplex (**Fig. 1*A***). The levels of complex V (analyzed using antibodies for the subunit ATP5A) were unchanged (**Fig. 1*A***). The defects observed in the supercomplexes and complex IV were rescued upon re-expression of COA7 with a C-terminal Flag epitope (+COA7^Flag^), indicating the complex IV defect observed was indeed due to the loss of COA7 (**Fig. 1*A***).

**Figure 1.**
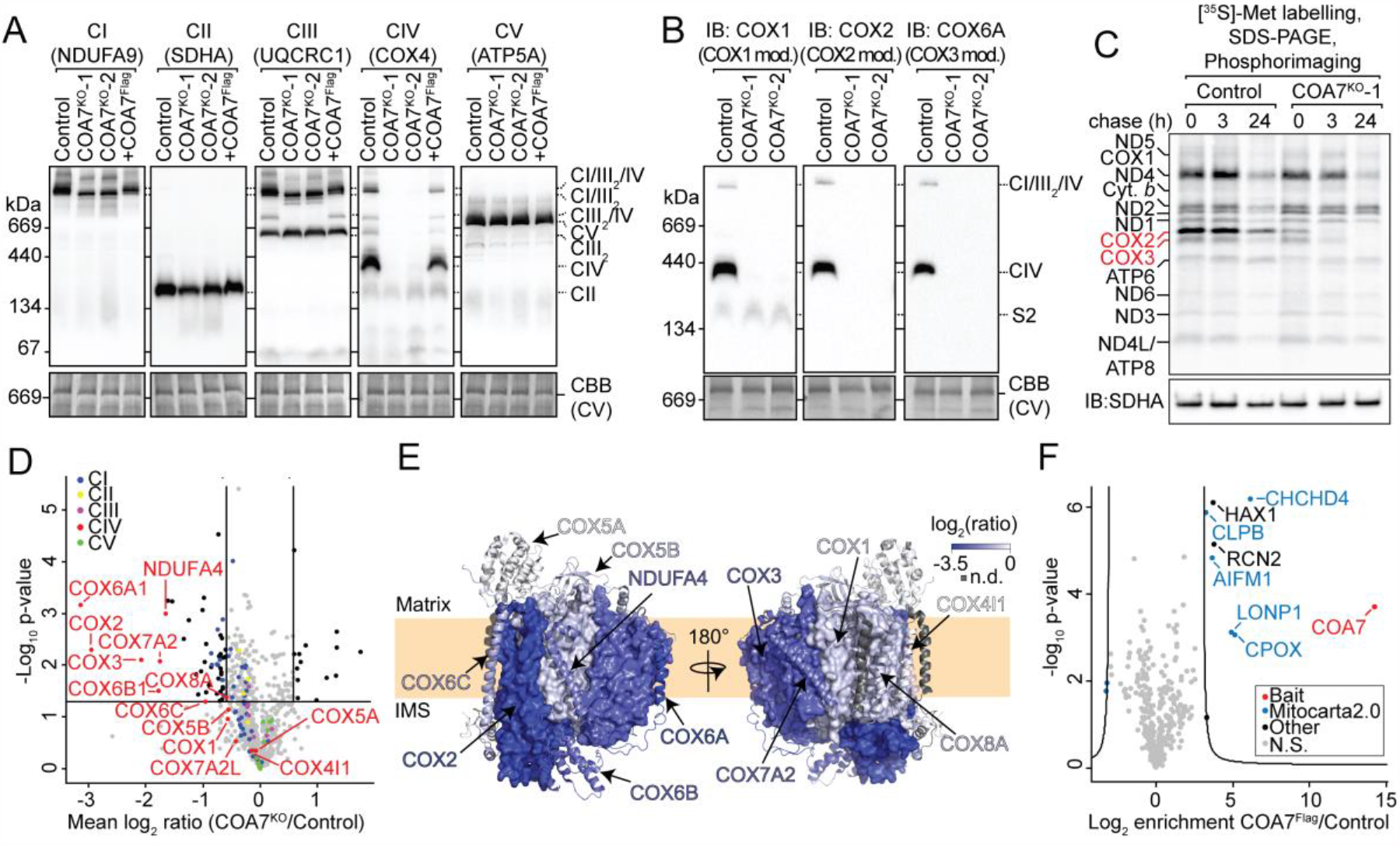
Loss of COA7 results in defects of respiratory chain enzymes. (*A*-*B*) Mitochondria isolated from control, COA7 knockout and rescued COA7 knockout cells (as indicated) were solubilized in 1% digitonin and subjected to BN-PAGE and analyzed using the antibodies shown. Coomassie brilliant blue (CBB) stained membranes are indicative of loading. (*C*) Cells were pulsed with [^35^S]-Met for 2 hours and chased for the indicated times. Isolated mitochondria were analyzed by SDS-PAGE and autoradiography. SDHA was used as a loading control. (*D*) Volcano plot showing proteins with altered abundance in COA7^KO^ mitochondria. Subunits of the different OXPHOS complexes are color coded. Complex IV subunits are labeled. The horizontal line indicates *p =* 0.05, vertical lines indicate a fold change of 1.5. (*E*) Topographical heatmap of SILAC log_2_(ratios) mapped onto the structure of complex IV (PDB 5Z62) (26). Proteins encoded by mtDNA are shown as surface renders, while nuclear encoded proteins shown as cartoon. Blue intensity is relative to the decrease in protein abundance, n.d. not detected. (*F*) Digitonin-solubilized mitochondria from control or COA7^KO^ + COA7^Flag^ cells were subjected to affinity enrichment using Flag agarose beads followed by label free quantitative proteomics. Significance was determined through a two-sided, two sample t-test using permutation-based FDR statistics. The curve represents the cutoff for FDR < 5% using an s0 parameter of 20.

To delineate the effect of COA7^KO^ on assembly of complex IV, we analyzed representative subunits of each of the three assembly modules: COX1 and COX2 and COX3 (detecting COX1, COX2 and COX6A, respectively) (**Fig. 1*B***). The analysis showed that residual COX1 was present in a faster migrating complex, termed the ‘S2’ intermediate (21), while COX2 and COX6A were not present in any observed intermediates (**Fig. 1*B***). To further investigate the stability of the core mtDNA-encoded proteins, we utilized [^35^S]-methionine pulse-chase labeling in the presence of anisomycin, an inhibitor of cytosolic translation (22). Control and COA7^KO^-1 cells were labeled for 2 hours with [^35^S]-Met before a chase without label and restoration of translation. Phosphorimage analysis SDS-PAGE showed that COX2 was highly labile in COA7^KO^ cells, with reduced labeling of this subunit at the zero-hour time-point and rapid loss over time, relative to the control cells (**Fig. 1*C***). COX3 also displayed increased turnover during the chase period in COA7^KO^ cells, compared with control mitochondria (**Fig. 1*C***). In comparison, translation and stability of COX1 was similar in both lines (**Fig. 1*C***). The levels of other mtDNA-encoded proteins did not appear to differ between control and COA7^KO^ mitochondria. These results point to a critical role for COA7 in complex IV assembly in steps that follow the formation of the COX1 module.

We next sought to understand the effect of COA7 loss on the mitoproteome using quantitative stable isotope labeling of amino acids in cell culture (SILAC) proteomics (23). In agreement with the pulse-chase analysis, a significant decrease in the abundance of COX2 and COX3 was observed (**Fig. 1*D*; Dataset S1**). In addition, the levels of a number of nuclear-encoded complex IV subunits (COX6A1, COX6B1, COX7A2, COX8A and NDUFA4), which are associated with the COX2 or COX3 modules or later complex IV assembly intermediates (4, 24, 25), were reduced in the absence of COA7 (**Fig. 1*D*; Dataset S1**). The visualization of the relative levels of complex IV subunits on the structure of complex IV using a topographical heatmap (PDB 5Z62) (26) shows that the COX1 module subunits (COX1, COX4 and COX5A) are largely unaffected by the loss of COA7, which is in stark contrast to the other mtDNA-encoded subunits COX2 and COX3, and their surrounding nuclear encoded proteins (**Fig. 1*E***). Given that we observe a reduced COX1 signal by BN-PAGE in COA7^KO^ mitochondria compared to control mitochondria, it is possible that COX1-containing assemblies subsequent to the S2 subcomplex are prone to aggregation once extracted from the membrane environment. This suggests that the COX1 module is unable to mature and integrate with other COX modules, resulting in turnover of the COX2 and COX3 modules. Taken together, these results indicate that COA7 plays a critical role in the assembly of complex IV at a step that involves association of the COX2 and COX3 modules onto the COX1 module.

We performed affinity-enrichment mass spectrometry to identify stable COA7 interacting proteins in an unbiased manner. Mitochondria isolated from control and COA7^KO^ + COA7^Flag^ rescue cells were solubilized in digitonin, incubated with anti-Flag beads and the bound proteins, eluted with Flag peptide, were subjected to label free quantitative (LFQ) proteomics analysis. COA7^Flag^ failed to enrich any complex IV structural proteins or assembly factors (**Fig. 1*F;* Dataset S2**). COA7^Flag^ however, did enrich a number of IMS proteins, including apoptosis-inducing factor 1 (AIFM1) and CHCHD4, which are required for disulfide bond formation in the intermembrane space (27). In addition, COA7 was able to enrich coproporphyrinogen oxidase (CPOX), an enzyme involved in heme biosynthesis (28), the mitochondrial protease LONP1 (29) and the disaggregase CLPB and the CLPB-interacting protein HAX1 (30) (**Fig. 1*F;* Dataset S2**). RCN2 an ER-luminal protein with potential localization in the mitochondrial IMS (31), was also identified in association with COA7^Flag^. These results suggest that unlike many assembly factors, COA7 does not form stable interactions with components of the respiratory chain.

### Structural characterization of COA7

To uncover further functional information, we sought to determine the molecular structure of COA7. Recombinant glutathione S-transferase (GST) fusions with COA7, pathogenic mutant proteins ^R39W^COA7, ^Y137C^COA7, and ^S149I^COA7 and additional variants (^Y137S^COA7, ^H119A^COA7 and ^M156A^COA7) were overexpressed and purified by affinity and size exclusion chromatography (SEC), yielding preparations of the GST-free proteins of approximate 90-95% purity as determined by SDS-PAGE, with intense bands at ∼25 kDa, corresponding to the molecular weight of monomeric COA7 (**Fig. S2**). The analytical SEC elution profile of the resulting purified COA7 showed a broad elution peak with an elution volume corresponding to a molecular weight of ∼55 kDa, indicating the possible presence of a dimeric species in solution (**Fig. S2**). The quaternary structure of COA7 was further investigated by analytical ultracentrifugation (AUC), using sedimentation velocity (SV) experiments. These data were fitted to a continuous size distribution model, which yielded sedimentation coefficients of 1.9 S and 3.5 S, demonstrating that the sample contained a mixture of monomeric and dimeric species (**Fig. S2, Table S2**), with the relative proportions of these species observed to vary with protein concentration, indicating concentration-dependent oligomerization behavior.

We determined the structure of COA7 by X-ray crystallography. The structure was solved and refined in space group *I*4_1_ to 2.4 Å resolution, with a single molecule of COA7 per asymmetric unit (**Fig. 2*A***). The final COA7 model includes residues Glu10-His218 and refinement of the model converged with residuals *R* = 20.5% and *R*_free_ = 25.5% (**Table S3**). COA7 is comprised of 11 α-helices, arranged as five helix-turn-helix (α/α) repeats (which vary in length from 30-36 residues) and an additional C-terminal helix (**Fig. 2*A***). Together, these form an elongated, right-handed super-helix, with a concave and a convex face. The COA7 sequence includes 13 cysteine residues (at positions 24, 28, 37, 62, 71, 95, 100, 111, 142, 150, 172, 179 and 187), with the structure showing five intramolecular disulfide bonds between residues Cys28-Cys37, Cys62-Cys71, Cys100-Cys111, Cys142-Cys150 and Cys179-Cys185 (**Fig. 2*B***), which bridge pairs of helices within each of the five α/α repeats. The ten Cys residues that participate in the intramolecular disulfide bonds are conserved in COA7 sequences across all metazoans however, the cysteine residues at positions 24, 95 and 172 (which are not observed to participate in disulfide bonds in this structure) are found in COA7 sequences from vertebrates only (**Fig. S1**) (10).

**Figure 2.**
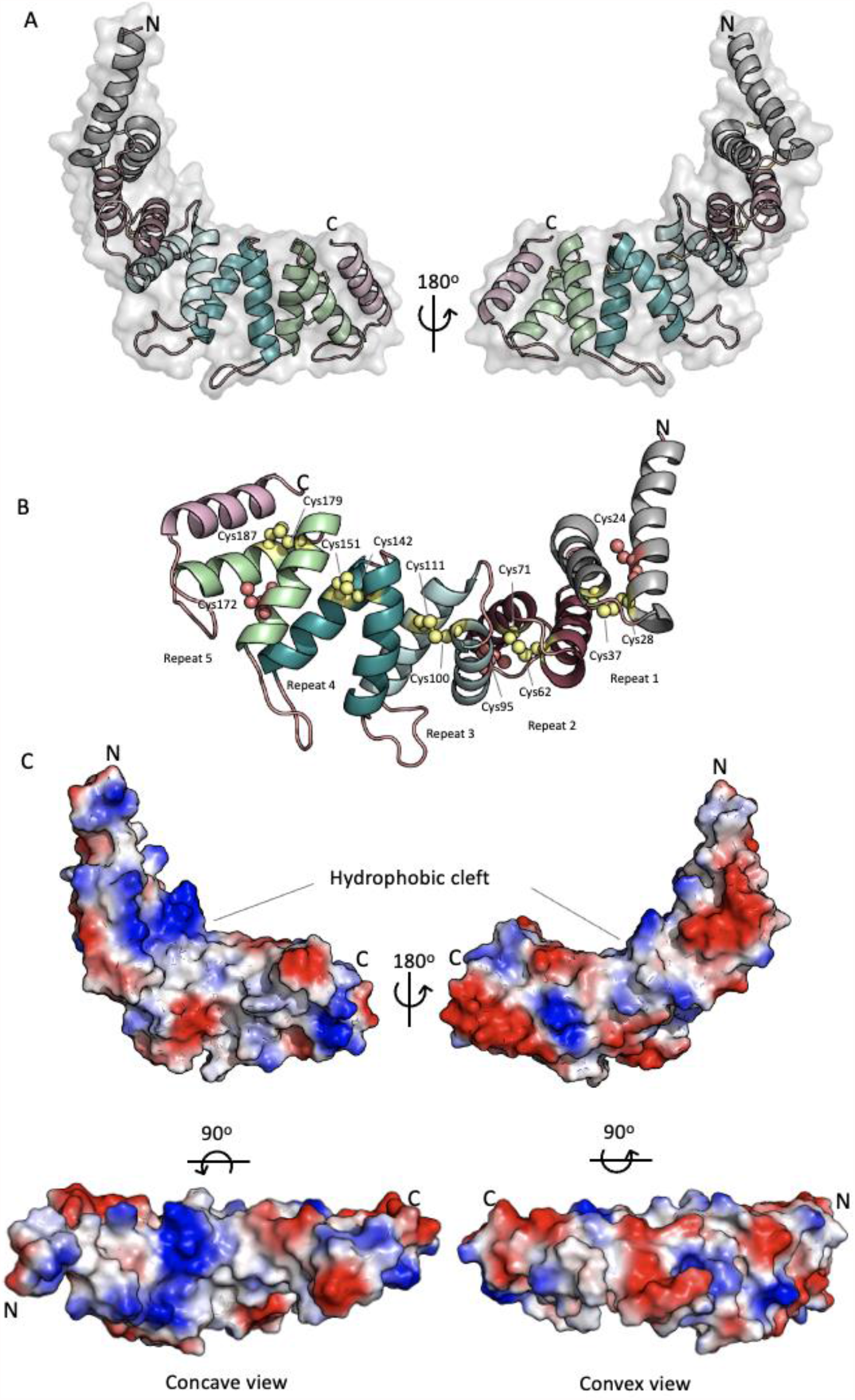
Structure of COA7. (*A*) Cartoon representation of the overall structure of COA7. Secondary structures are represented as cartoons with SEL1 repeats colored in gray, raspberry, cyan, teal and green. The transparent molecular surface is represented in gray (*B*) Cysteine residues (labeled) involved in intramolecular disulfide bonds are shown as yellow spheres, and free cysteine residues are shown as salmon spheres. (*C*) The molecular surface of COA7 is colored according to the electrostatic potentials (red, negatively charged; blue, positively charged; white, uncharged). The top images show the side view of the COA7 structure. The left and right bottom images show concave and convex views, respectively.

A search against the PDB (32) revealed that COA7 shares significant structural similarity with the <u>H</u>elicobacter <u>c</u>ysteine-rich <u>p</u>roteins B and C (HcpB, (PDB 1KLX) and HcpC (PDB 1OUV), respectively) (33, 34) from *Helicobacter pylori* (**Fig. S3**). Hcp proteins are composed of α/α repeats and belong to the SLR protein family. The presence of disulfide bonds within the repeats for both the Hcps and COA7 is unusual for this protein family (13, 33). Indeed, to our knowledge, COA7 is the first structurally characterized human SLR protein possessing disulfide-bridged α/α repeats.

The outer convex surface of the COA7 molecule is highly negatively charged due to the presence of a number of Glu and Asp residues (located within repeats 1, 2 and the linkers between repeats 3, 4 and 4, 5 and the C-terminal helix). In addition, the presence of lysine residues 49, 56, 59, 73, 82, 105 and 106 on the inner concave surface of the COA7 molecule, creates a positively charged surface in this region (**Fig. 2*C***). The residues that comprise both of these charged surfaces are largely conserved in COA7 proteins (**Fig. S1**). Also on the concave surface is a hydrophobic cleft that results from clustering of several hydrophobic residues (Ile108, Ala109, Gly144, Gly145, Leu181 and Gly182) (**Fig. 2*C***). Interestingly, the application of crystallographic symmetry operators reveals that in the crystal, this cleft is occupied by the N-terminal helix (residues Glu10-His30) of a neighboring molecule of COA7 (**Fig. S4**), such that the crystal is composed of an infinite network of these protein-protein interactions (**Fig. S4**). This interaction is mediated by hydrogen bonds and electrostatic interactions, with a total buried surface area of 1614 Å^2^, representing ∼7% of the surface areas of the two molecules combined (32). Given the limited extent of this interaction, it is unlikely to represent a persistent oligomeric structure in solution (35), but correlates with the concentration-dependent oligomerization behavior observed by AUC. However, this contact may indicate a molecular mechanism for an interaction between COA7 and protein partners or ligands in the mitochondrial IMS. Similar direct protein-protein interactions have been previously proposed for other TPR proteins, including HcpC (34), the Hsp70/Hsp90 organizing protein (Hop) (PDB 1ELR) (36, 37) and the peroxisomal targeting signal-1 PEX5 (PDB 1FCH) (38). In the case of the related HcpC structure, an interaction between the C-terminus of one molecule and the hydrophobic pocket of a neighboring molecule facilitates the formation of a similar, extended protein network in the crystal. This was proposed to represent a mechanism for protein-protein interactions in the cell (34).

### COA7 pathogenic mutations

Pathogenic mutations in the *COA7* gene producing the mutant variants ^D6G^COA7, ^R39W^COA7, ^Y137C^COA7 and ^S149I^COA7 have been identified in patients exhibiting neurological symptoms of peripheral neuropathy, ataxia and leukoencephalopathy (18, 19). Analysis of the crystal structure of COA7 reveals that residue Arg39 is located on the C-terminal helix of repeat 1 and is exposed on the outer surface of the COA7 molecule (**Fig. 3*A***). Residues Tyr137 and Ser149 are located in the ‘middle’ of the structure, on repeat 4, and participate in hydrogen bonding interactions with residues Glu101 and His112, respectively which are both found on repeat 3 (**Fig. 3*A***). Residue Asp6 is at the N-terminus of the molecule and could not be resolved in the crystal structure.

**Figure 3.**
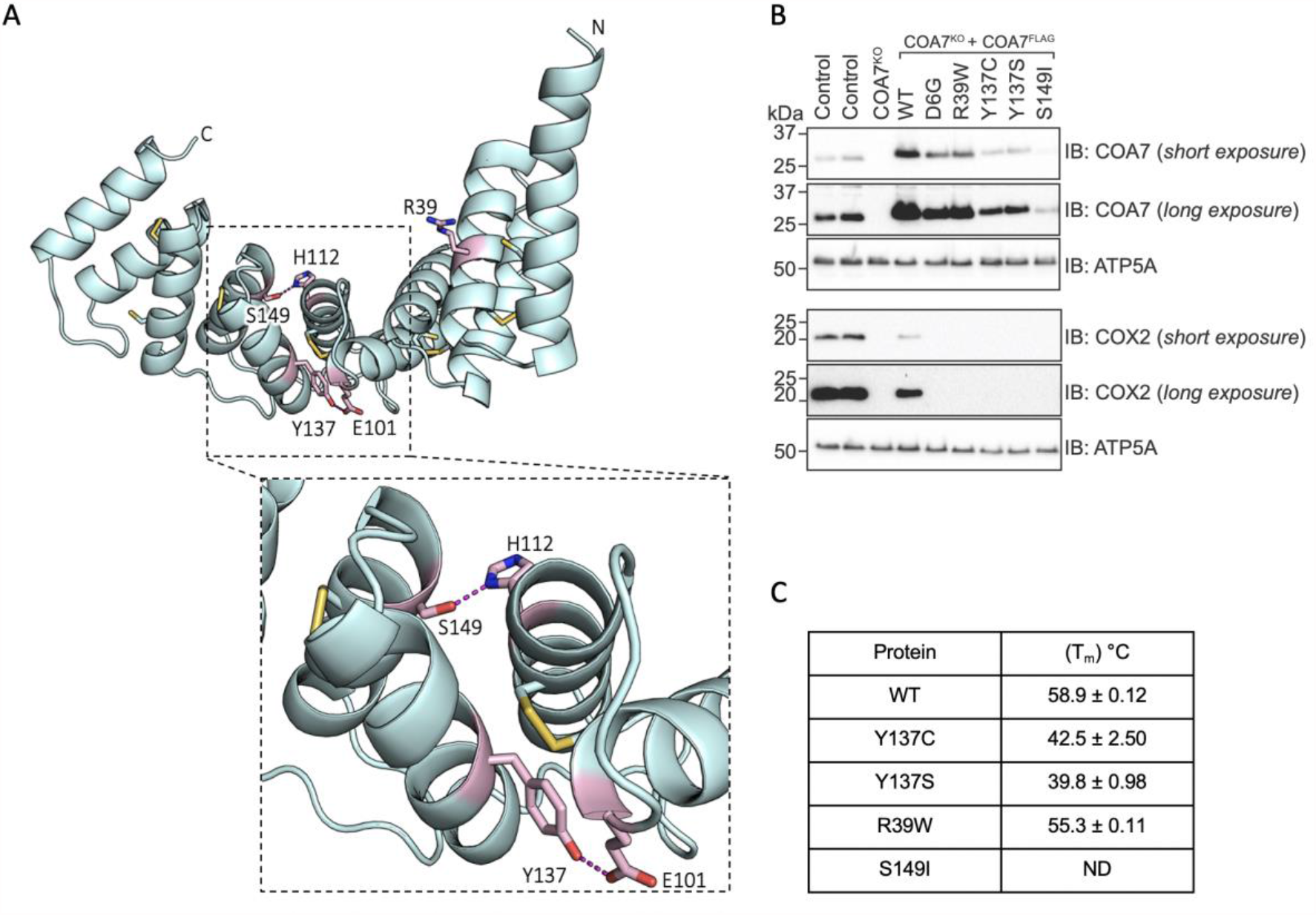
COA7 pathogenic mutations. (*A*) The structure of COA7. Patient mutations are shown as pink sticks and hydrogen bonds between residues (labeled) shown as dashed lines. Inset: closer view of the interactions formed by the His112 and Y137 residues. (*B*) Mitochondria isolated from control, COA7^KO^ cells and COA7^KO^ cells expressing COA7^Flag^ and variants as indicated were subjected to SDS-PAGE and immunoblotting with antibodies as shown. ATP5A serves as a loading control. (*C*) Melting temperatures of the COA7 and variant proteins. These values were determined by differential scanning fluorimetry from purified protein samples.

In order to investigate the effects of the pathogenic mutations identified in patients, we generated doxycycline-inducible retroviral plasmids encoding COA7 mutant proteins for stable expression in COA7^KO^ cells. Expression of COA7 with a Flag epitope tag was able to rescue COX2 levels (although not to control levels) in the COA7^KO^ line (**Fig. 3*B***). However, upon expression of the patient derived COA7 variants (^D6G^COA7, ^R39W^COA7, ^Y137C^COA7 and ^S149I^COA7), the levels of COX2 were not able to be restored to the same extent, indicating that these variants were less active in complex IV assembly, relative to COA7 (**Fig. 3*B***). Expression of the ^Y137S^COA7 protein was also tested to determine if an amino acid substitution of similar geometry to the Y137C patient-identified mutation, but unable to form disulfide bridges, was functional. The Y137S mutant was not able to rescue the defect and showed a similar impact as the Y137C mutation (**Fig. 3*B***).

Recombinant, purified forms of ^R39W^COA7, ^Y137C^COA7, ^S149I^COA7 and ^Y137S^COA7 were prepared and subjected to differential scanning fluorimetry (DSF, **Fig. 3*C***) to measure their thermal stabilities (39). Reduced protein stabilities, relative to COA7 were observed for all variant proteins, consistent with the observed decreased expression in mitochondria (**Fig. 3*B***). In particular, the melting temperatures of the ^Y137C^COA7 and ^Y137S^COA7 proteins were 16-19°C lower than COA7. The thermal transition for purified ^S149I^COA7 could not be observed (**Fig. 3*C***), indicating the protein was most likely misfolded. Analysis of the quaternary structure of ^Y137C^COA7 by analytical ultracentrifugation showed evidence of concentration dependent dimerization (as observed for COA7), in addition to the presence of higher order oligomeric states indicative of aggregation. These observations may explain the reported aggregation and mislocalization of the ^Y137C^COA7 protein in cells (10) (**Fig. S5, Table S2**).

A recent analysis of the COA7 and ^Y137C^COA7 proteins *via* molecular modeling and simulation studies, proposed that the presence of residue Cys137, may lead to the disruption of the Cys100-Cys111 disulfide bond and the dynamic formation of alternative disulfide bonds between the introduced Cys137 and residues Cys100 and Cys111 (10). The structure reported here shows that the formation of an alternative Cys137-Cys111 disulfide bond would be possible without major structural rearrangement. However, the distance between the thiolate groups of the introduced Cys137 and residue Cys100 would be too long (3.9 Å) in the absence of a major structural reorganization (**Fig. S6**). Indeed, the Y137S mutation, which cannot form an alternative disulfide bond, resembled the Y137C mutation with respect to its melting temperature (**Fig. 3*C***) and inability to rescue the COA7^KO^ phenotype (**Fig. 3*B***). These findings lead us to conclude that a rearrangement of the disulfide bonding network of COA7 upon mutation is unlikely to be a factor in the pathogenic effects of this variant form of the protein.

The COA7 S149I pathogenic mutation is predicted to eliminate the hydrogen bond with residue His112 and the larger Ile sidechain may also disrupt the helical packing between repeats 3 and 4 (**Fig. 3*A***). Given that ^S149I^COA7 was likely misfolded and that both the Y137C and S149I mutations are located in the ‘centre’ of the crescent-shaped structure (**Fig. 3*A***), this suggests that hydrogen bonds and other intramolecular interactions in this region may provide rigidity within the structure, most likely scaffolding one half of the assembly relative to the other. Loss of these interactions lead to instability of the proteins and decreased activities in terms of complex IV biogenesis. Finally, in the case of R39W variant, residue Arg39 is exposed on the outer surface of the COA7 molecule. The introduction of the R39W mutation is predicted to alter the charge in this region of the protein surface. The DSF analysis of the ^R39W^COA7 variant protein showed it to be well folded (**Fig. 4*C***). The pathogenesis of this mutation may therefore lie in altered interactions of this protein with as yet unidentified partners/substrates, associated with the change in the electrostatic structure of its surface.

**Figure 4.**
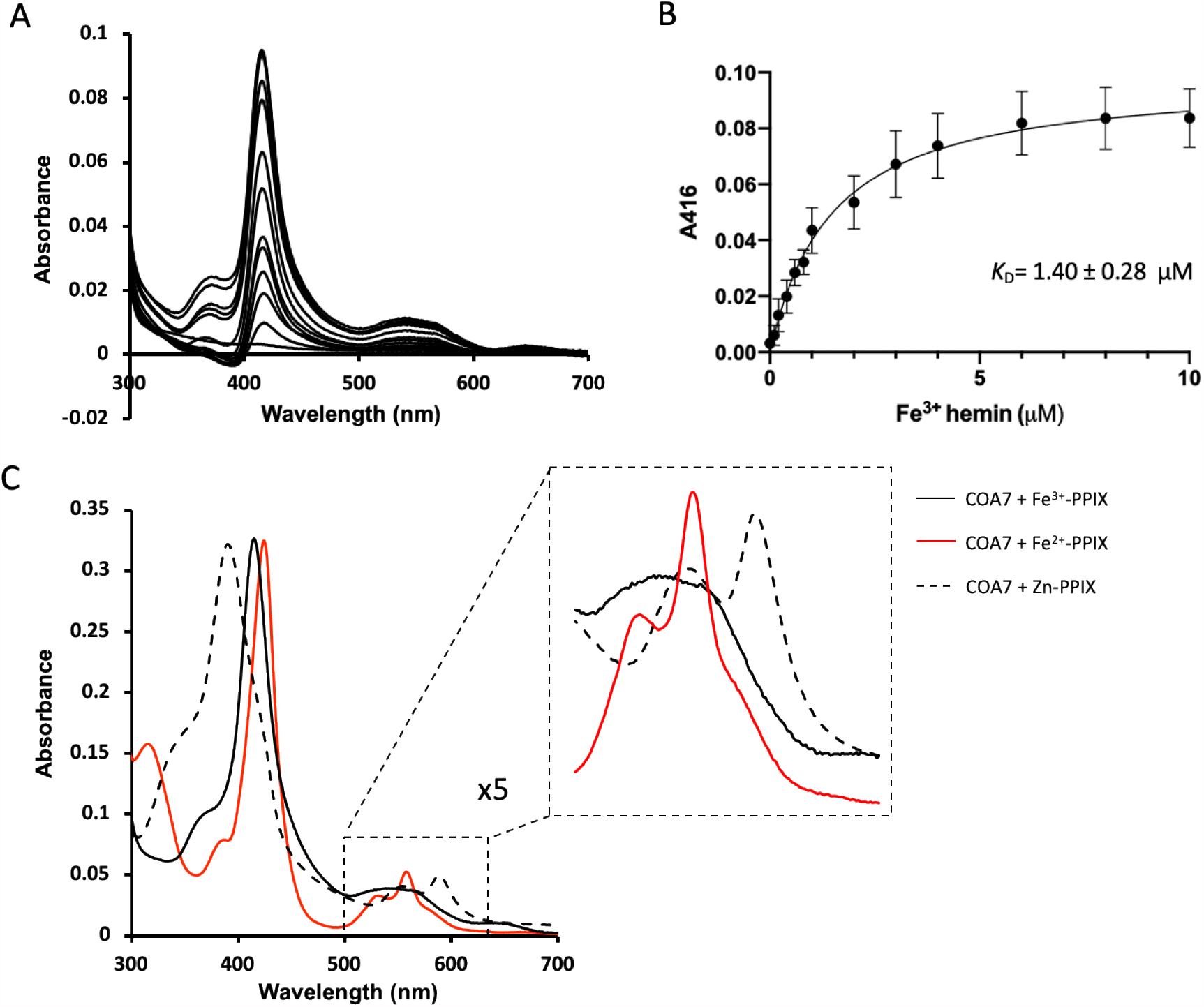
Heme binding to COA7. (*A*) Difference absorption spectra of COA7 (1 μM) with increasing concentrations of Fe^3+^-PPIX (from 0 to 10 μM) as indicated. (*B*) The binding curve was fitted to the data from (A) using an equation describing a single binding site (Y = Bmax * X/*K*_D_ + X) with GraphPad Prism. (*C*) UV-Vis absorption spectra of COA7 in the presence of one molar equivalent of Fe^3+^-PPIX (black); Fe^2+^-PPIX (red), and Zn^2+^-PPIX (dotted, black).

### COA7 binds heme with micromolar affinity

A key step in the biogenesis of complex IV is insertion of copper into COX2 from the IMS, which involves a suite of assembly factors (6). Loss of function mutations in these assembly factors blocks complex IV assembly at the level of COX2 maturation, similar to that observed here for COA7 (21). Furthermore, soluble high affinity Cu(I)-binding proteins (copper metallochaperones) such as ATOX1 (40), COX17 (41) and the recently identified Cu(I)-binding protein, COA6 (21) all show CysX_n_Cys sequence motifs, bind Cu(I) and function in copper delivery to protein partners within the cell. Due to the presence of multiple Cys residues within the COA7 sequence, we sought to determine whether COA7 binds copper (specifically Cu(I)) using a competition assay with the [Cu^I^Bcs_2_]^3−^ complex (21, 42). Cu(I)-binding to the COA7 was not detected using this assay, however Cu(I) binding was observed for the recombinant ATOX1 protein, which was included as a positive control (**Fig. S7**).

Given these findings, along with our inability to detect an association between COA7 and other copper chaperones, we investigated whether COA7 might bind heme, the other cofactor required for complex IV biogenesis and activity. Complex IV binds heme *a* and heme *a*_*3*_, which are essential for function. Heme *a* is important for the stability of COX1, while insertion of heme *a*_*3*,_ which has a farnesyl tail and sits between COX1 and COX2, is required for assembly and stabilization of COX2 (5, 6). We therefore tested whether COA7 might be involved in heme incorporation into complex IV by measuring whether the purified protein was able to bind heme in solution using ultra violet-visible (UV-Vis) spectrophotometry (43, 44). Upon titration of COA7 with hemin, difference spectra revealed a Soret band at 416 nm and a broad feature between 500-600 nm (**Fig. 4*A***), which increased in intensity with the addition of increasing concentrations of hemin. Fitting of these titration data yielded *K*_D(heme)_= 1.40 ± 0.28 μM (**Fig. 4*B***). This value is similar to those measured for other heme binding proteins. For instance, the HemQ (chlorite dismutase-like protein) from *Listeria monocytogenes*, which is involved in heme acquisition by the bacterium binds heme with *K*_D(heme)_= 16.2 μM (45). The cytoplasmic heme binding protein Phus from *Pseudomonas aeruginosa* binds heme with *K*_D(heme)_= 29 μM (46).

Incubation of purified COA7 with one molar equivalent of hemin and reduction with sodium dithionite, yielded UV-Vis spectra indicative of heme-bound COA7 (**Fig. 4*C***). These spectra were very similar to the those reported for the ferric and ferrous states, respectively of cytochrome *b*_562_ from *E. coli* (47), which shows axial ligation of the heme iron by His and Met residues. Incubation of COA7 with Zn^2+^-PPIX also yielded a spectrum very similar to that of Zn^2+^-PPIX-loaded cytochrome *b*_562_ (**Fig. 4*C***) (47). The incubation of COA7 with excess heme, followed by removal of unbound cofactor, yielded a COA7-heme complex with a stoichiometry, as determined by inductively-coupled plasma mass spectrometry (ICP-MS), of 1.06 ± 0.03 iron atoms (heme cofactors) per protein molecule (**Fig. S8**). COA7 could not be co-purified with PPIX using the same approach (**Fig. S8**).

These experiments indicate specific heme binding to COA7, with spectral evidence for axial ligation to the iron through His and Met ligands. The only TPR protein characterized to bind heme is the HusA protein from *Porphyromonas gingivalis*, with a structure of four α/α repeats, which in contrast to COA7, exhibited iron-independent heme binding (that is, no direct coordination from the protein to the heme iron) (48). Aside from the presence of the TPRs, the HusA structure bears little similarity to that of COA7 in that it shows a relatively ‘closed’ structure compared with COA7 and does not include disulfide bonds. Examination of the structure of COA7 reveals a possible heme binding site located between repeats 3 and 4, with residues His119 and Met156 in comparable positions to those that act as iron ligands in cytochrome *b*_562_ (47) (**Fig. S9*A***). We therefore generated single and double mutant proteins (^H119A^COA7, ^M156A^COA7 and ^H119A/M156A^COA7) to probe the contributions of these residues to heme binding.

Titration of the ^H119A/M156A^COA7 variant with hemin, yielded no detectable binding (**Fig. S10*A*-*C***) and the UV-Vis spectra of the variant protein following incubation with one equivalent of hemin were significantly different from that observed for COA7 (both in the presence and absence of sodium dithionite; **Fig. S11**). Finally, we incubated the heme-loaded COA7, ^H119A^COA7 and ^M156A^COA7 proteins with sodium cyanide and compared the UV-Vis spectra of the proteins measured before and after incubation (**Fig. S10*D-F***). The spectrum of the heme-COA7 protein showed no change on incubation with CN^-^. In contrast, for both the heme-^H119A^COA7 and heme-^M156A^COA7 variants, shifts in the positions of the Soret bands and spectral changes in the region between 550-650 nm were observed, indicating a change in iron coordination (therefore ligand substitution) on addition of CN^-^. These data taken together, indicate that heme binding to COA7 is mediated by axial ligation of residues His119 and Met156 to the heme iron. A superposition of the COA7 structure with that of cytochrome *b*_562_ (PDB 256B; **Fig. S9*B***,***C***) (49), shows that heme binding in this position, comparable to that seen in the structure of cytochrome *b*_562_ (49), would be possible upon slight adjustments to the positioning of the surrounding helices. These data indicate that *in vitro* COA7 binds heme specifically through axial ligation of the central iron atom with His119 and Met156 protein ligands, which are conserved in the majority of COA7 sequences (**Fig. S1**).

## Conclusion

Despite the importance of the assembly of complex IV in health and mitochondrial disease, we have an incomplete understanding of the molecular basis of the biogenesis of this enzyme owing to a lack of knowledge about the identities, structures and roles of all crucial assembly factors (6). The present study shows that deletion of COA7 resulted in a severe complex IV assembly defect. Crucially, the absence of COA7 causes specific defects in the maturation of the COX2 and COX3 modules, resulting in their degradation and therefore halting complex IV biogenesis. This suggests that the COX1 module may be unable to fully mature and integrate with the COX2 and COX3 modules.

The high-resolution crystal structure of the human COA7 protein shows the protein is composed of five SLRs, tethered by disulfide bonds. The overall fold of COA7 is similar to other TPRs and SLRs, proposed to function in protein-protein interactions due to their extensive solvent accessible surfaces (13, 15-17). Fascinatingly, recombinant COA7 was found to bind heme in solution *via* axial His/Met coordination to the heme iron, despite showing no structural similarity to well-characterized heme binding folds. A role for COA7 in chaperoning heme in the IMS for insertion into complex IV would be consistent with the observed ‘blockage’ of complex IV assembly at the COX1 module in the absence of the assembly factor.

Heme is a highly reactive and inherently toxic compound. As a cofactor, heme *b* is required for the activities of a plethora of hemeoproteins, present in virtually every subcellular compartment throughout the eukaryotic cell. It is also a precursor for the synthesis of other heme types important for eukaryotic physiology, including hemes *a, c* and *o*. All of these processes require that heme is trafficked from its site of synthesis in the mitochondrial matrix to heme-dependent proteins throughout the cell (28). For complex IV biogenesis, heme *b* is converted to heme *a* through the sequential activities of the enzymes COX10 (which catalyzes the synthesis of heme *o*) and COX15, which have active sites facing the matrix and the IMS, respectively (28, 50). To date, no information is available on how heme *o* is transported across the IMM to the COX15 active site. Indeed, the aqueous insolubility and instability of hemes in general preclude the existence of ‘free’ (uncomplexed) forms of this cofactor, although low concentrations (<2.5 - 40 nM) of labile heme pools exist in different cellular compartments (51). Given the data presented here, it is tantalizing to suggest then that COA7 provides a chaperoning role for heme *o* and/or heme *a* in the IMS. If so, this would certainly provide a missing link for heme handling in mitochondria.

## Online Methods

### Cell culture and genome editing

Human epithelial kidney cells containing the SV40 T-antigen (HEK293T) were purchased from the ATCC and a previously established clonal line was used (52, 53). Cells were cultured in DMEM supplemented with 10% (v/v) fetal bovine serum (FBS), 1 × penicillin/ streptomycin (Sigma-Aldrich; P4458), 1 × Glutamax (Life Technologies; 35050061) and 50 μg/mL uridine (Sigma-Aldrich; U3750). Cells were grown at 37°C with a humidified 5% CO_2_ atmosphere. Galactose containing DMEM was prepared from glucose-free DMEM (Life Technologies; 11966-025) supplemented with 10% (v/v) dialysed FBS (GE Healthcare; SH30079.03), 25mM D-galactose (Sigma-Aldrich; G0750), 1 × penicillin/ streptomycin, 1 × Glutamax, 1 × sodium pyruvate and 50 μg/mL uridine.

For SILAC culture, cells were grown as previously described (52). Briefly, cells were grown in DMEM without lysine/arginine, supplemented with 10% (v/v) dialyzed FBS, 1 × penicillin/ streptomycin, 1 × Glutamax, 1 × sodium pyruvate and 50 μg/mL uridine and either ‘heavy’ amino acids (^13^C_6_^15^N_4_-Arginine; Cambridge Isotope Labs Inc; CNLM-539-H-1, ^13^C_6_^15^N_2_-Lysine; Silantes; 211604102) or ‘light’ amino acids (arginine; Sigma-Aldrich; A5131, lysine; Sigma-Aldrich; L5626).

Gene editing was performed using the pSp-CAS9(BB)-2A-GFP (PX458) CRISPR/Cas9 construct (a gift from F. Zhang; Addgene plasmid 48138) (20). The guide RNA was designed to target the 3’ end of the first exon with the sequence 5’-CTGCTACCACGAGAAGGACC-3’. Indel sequencing of the genomic locus where the CRISPR/Cas9 was targeted demonstrated that COA7^KO^-1 contained the following mutations c.[93_99del];[101_102insC] leading to p.[E31Dfs*25; D35Gfs*17]. Analysis of COA7^KO^-2 demonstrated this cell line contained the mutations c.[100_101 insTAATATCTTTGTGTTTACAGTCAAATTAATTCCAATTATCTCTCTAACAGCCTT GTATCGTATATGCAAATATGAAGGAATCATGGGAAATAGGCCCTCACATGTGAGCAAAAGGCCAGCAAAAGGCCAGGAACCGTAAAAAGGCCGCGTTGCTGGCGTTTTTCCATAGGCTCCGCCCCCCTGACGAGCATCACAAAAATCGACGCTCAAGTCAGAGGTGGCGAAACCCGACAGGACTATAAAGATACCAGGCGTTTCCCCCTGGAAGCTCCCTCGTGCGCTCTCCTGTTCCGACCCTGCCGCTTACCGGATACCTGTCCGC; 101_102ins AATATCTTTGTGTTTACAGTCAAATTAATTCCAATTATCTCTCTAACAGCCTTGTATCGTATATGCAAATATGAAGGAATCATGGGAA ATAGGCCCTCACATGTGAGCAAAAGGCCAGCAAAAGGCCAGGAACCGTAAAAAGGCCGCGTTGCTGGCGTTTTTCCATAGGCTCCGCCCCCCTGACGAGCATCACAAAAATCGACGCTCAAGTCAGAGGTGGCGAAACCCGACAGGACTATAAAGATACCAGGCGTTTCCCCCTGGAAGCTCCCTCGTGCGCTCTCCTGTTCCGACCCTGCCGCTTACCGGATACCTGTCCGCC] leading to p.[P34Lfs*9;D35Ifs*8].

### Transfection and stable cell line generation

Cells were transfected using Lipofectamine LTX (Life Technologies; 15338100) according to manufacturer’s instructions. Stable, inducible expression was achieved by transfecting cells with pLVX-TetOne-Puro (Clonetech; 631849) encoding the wildtype COA7 cDNA. For mutational analysis in COA7^KO^ cells, wildtype or mutant COA7^Flag^ was cloned into pRetroX-TetOne-Puro (Clonetech; 634309), downstream of a GFP-T2A sequence to facilitate sorting of COA7-expressing cells (54). The COA7 cDNA was amplified from a HEK293T cDNA library derived from the clonal HEK293T cells. Mutant COA7 cDNA was generated using mutagenic primers to generate PCR fragments that incorporate the desired mutation, which were then subsequently assembled using Gibson Assembly (NEB). All plasmids were sequence verified by Micromon (Monash University).

Lentiviral supernatants were prepared by co-transfecting HEK293T cells with pLVX-TetOne-Puro plasmid encoding the cDNA of interest together with pVSV-G and pSPAX2 helper vectors. Retroviral supernatants were prepared by co-transfecting HEK293T cells with pRetroX-TetOne-Puro plasmid encoding the cDNA of interest together with pVSV-G and Gag-Pol helper vectors. Supernatants were collected after 48h and filtered through a 0.45µm low protein binding filter (Merck). COA7^KO^-1 cells were then infected with the lenti/retroviral supernatant with the addition of 8 µg/mL polybrene for stable and inducible expression of COA7^Flag^. Transduced cells were selected by the addition of 2 µg/mL puromycin until control cells died. Cells transduced with pRetroX-TetOne-Puro COA7^Flag^ variants were selected by fluorescence-activated cell sorting (FACS), after expression of GFP.

### mtDNA subunit labeling

Radiolabeling of mtDNA-encoded proteins was performed using [^35^S]-Methionine/Cysteine (Perkin Elmer) in the presence of 10 μg/mL Anisomycin (Sigma-Aldrich; A9789) to block cytosolic translation as previously described (22). Labeling was performed for 2 hours, followed by a chase with regular DMEM for 0, 3 and 24 hours. Isolated mitochondria were then analyzed by Tris-Tricine SDS-PAGE and phosphorimagining digital radiography (GE Healthcare).

### Mitochondrial isolation, whole cell lysate preparation, gel electrophoresis and western blotting

Mitochondria were isolated by differential centrifugation as previously described (53, 55). Protein concentration was determined by bicinchoninic acid assay (BCA; Thermo Fisher Scientific; 23223). Mitochondria were used immediately or aliquoted and frozen at −80°C until required.

Tris-tricine SDS-PAGE and BN-PAGE were performed as previously described (56-58). For tris-tricine SDS-PAGE, mitochondria were solubilized in 1× LDS Sample Buffer (Thermo Fisher Scientific; NP0008) supplemented with 100 mM DTT and run on 10% acrylamide gels. Radiolabeling samples were analyzed using continuous 10-16% Tris-tricine SDS-PAGE gels. For BN-PAGE, mitochondria were solubilized in detergent buffer (20 mM Bis-Tris pH 7.0, 50 mM NaCl, 10% (w/v) glycerol) containing 1% digitonin for 10 minutes on ice, followed by centrifugation at 16,000 *g* for 10 minutes. The clarified supernatant containing mitochondrial complexes was then transferred to a clean tube containing one-tenth of the volume of 10x BN-PAGE loading dye (5% (w/v) Coomassie blue G250, 500 mM ε-amino n-caproic acid, 100 mM Bis-Tris pH 7.0) before loading onto the gel. Following electrophoresis, transfer onto PVDF membrane (Merck; IPVH00010) was performed using a Novex Semi-Dry Blotter (Thermo Fisher) or an Invitrogen Power Blotter System (ThermoFisher) according to manufacturer’s instructions.

Primary antibodies used in this study were α-COA7 (25361-1-AP; Proteintech), α-NDUFA9 (in-house), α-SDHA (ab14715; Abcam), α-UQCRC1 (459140; ThermoFisher), α-COX1 (459600; ThermoFisher), α-COX2 (A-6404; ThermoFisher), α-COX4 (ab110261; Abcam), α-COX6A (11460-1-AP; Proteintech) and α-ATP5A (ab14748; Abcam). Anti-mouse (Sigma-Aldrich; A9044) or anti-rabbit (Sigma-Aldrich; A0545) horseradish peroxidase-conjugated secondary antibodies were used at a dilution of 1:10,000. Clarity western ECL chemiluminescent substrate (BioRad; 1705061) was used for detection on a BioRad ChemiDoc XRS+ imaging system according to the manufacturer’s instructions.

### SILAC proteomics

SILAC proteomics was performed as described previously (52). Briefly, equal amounts of differently labeled ‘heavy’ and ‘light’ cells were mixed and mitochondria were subsequently isolated. Analysis consisted of two light control/heavy COA7^KO^ pairs and a label switched heavy control/light COA7^KO^ sample. For sample preparation, 100 µg of isolated mitochondria were solubilized in 1% (w/v) SDC, 100 mM Tris-Cl pH 8.1, 40 mM chloroacetamide and 10 mM TCEP prior to vortexing and heating for 5 minutes at 99°C with 1500 rpm shaking. Samples were then sonicated for 15 minutes in a room temperature water bath prior to the addition of 1 μg trypsin (Promega; V5113) and incubation overnight at 37°C. The supernatant was then transferred to 3x 14G 3M™Empore™ SDB-RPS stage tips (59). Ethyl acetate (99%) and 1% TFA was added to the tip before centrifugation at 3000 *g* at room temperature as described (59). Stage tips were washed first with 99% ethyl acetate and 1% TFA and then with ethyl acetate supplemented with 0.2% TFA. Samples were eluted in 80% acetonitrile (ACN) and 1% NH_4_OH and acidified to a final concentration of 1% TFA prior to drying in a SpeedVac.

### Affinity enrichment mass spectrometry

Affinity enrichment was performed as previously described (60). Mitochondria from control or COA7^KO^ + COA7^Flag^ expressing cells were isolated and solubilized in 1% (w/v) digitonin solubilization buffer (20mM Bis-Tris (pH 7.0), 50 mM NaCl, 10% (v/v) glycerol). Following clarification of lysates by centrifugation (16,000 *g*, 10 mins, 4°C), supernatants were applied to Flag affinity gel (Sigma) and incubated for 2 hours at 4°C with gentle rotation. Affinity gel was washed with 0.1% digitonin in solubilization buffer. Proteins were eluted using 150 μg/mL Flag peptide in solubilization buffer with 0.1% (w/v) digitonin. Elutions were mixed with 5 volumes of ice-cold acetone and incubated at -20°C overnight. Precipitated proteins were then pelleted by centrifugation (16,000 *g*, 10 mins, 4°C) and subsequently solubilized in 50 mM ammonium bicarbonate and 8M urea. This was then sonicated in a water bath at room temperature for 15 min before addition of Tris(2-carboxyethyl)phosphine hydrochloride (TCEP) to a final concentration of 5 mM and chloroacetamide (Sigma Aldrich) to a final concentration of 50 mM and the sample was incubated at 37 °C for 30 mins while shaking. The sample was then treated with ammonium bicarbonate to dilute the urea to a final concentration of 2 M. Trypsin (1 µg; Promega) was added to the sample before incubation overnight at 37°C. After this, the sample was acidified with 10% trifluoroacetic acid (TFA) to a final concentration of 1%. Stage tips were generated with two plugs of 3 M™ Empore™ SDB-XC Extraction Disks (Fisher Scientific, Pittsburgh, PA) that were activated with 100% acetonitrile (ACN) via centrifugation. All spins were performed at 1800 *g*. The tips were washed with 0.1% TFA, 2% ACN three times. The sample was added to the stage tip and eluted with 80% ACN and 0.1% TFA. The eluates were subsequently dried down using a SpeedVac.

### Mass-spectrometry and Data Analysis

Peptides from the above procedures were reconstituted in 2% ACN (v/v), 0.1% TFA (v/v) and transferred to autosampler vials for analysis by online nano-HPLC/electrospray ionization-MS/MS on either a ThermoFisher Scientific Orbitrap Q Exactive Plus (SILAC) or Orbitrap Elite Hybrid Ion-Trap (AE-MS) instrument as follows:

For SILAC mitochondria, data were acquired on an Orbitrap Q Exactive Plus instrument connected to an Ultimate 3000 HPLC. Peptides were loaded onto a trap column (Dionex-C18 trap column 75 μm × 2 cm, 3 μm, particle size, 100 Å pore size; ThermoFisher Scientific) at 5 μl/min before switching the pre-column in line with the analytical column (Dionex-C18 analytical column 75 μm × 50 cm, 2 μm particle size, 100 Å pore size; ThermoFisher Scientific). The separation of peptides was performed at 300 nl/min using a 125 min non-linear ACN gradient of buffer A [0.1% formic acid, 2% ACN, 5% DMSO] and buffer B [0.1% formic acid in ACN, 5% DMSO]. The mass spectrometer was operated in positive-ionization mode with spray voltage set at 1.9 kV and source temperature at 275 °C. Lockmass of 401.92272 from DMSO was used. Data were collected using the Data Dependent Acquisition using m/z 375–1400 m/z scan range at 70,000 m/z resolution with AGC target of 3e6. HCD for MS/MS was performed on the top 15 most intense ions with charges ≥2. Fragment ion spectra were acquired in Orbitrap at 17,500 m/z resolution, AGC target of 5e4 and maximum injection time of 50ms. Dynamic exclusion with of 30 seconds was applied for repeated precursors.

AE-MS was acquired on an Orbitrap Elite Hybrid Ion-Trap instrument connected to an Ultimate 3000 HPLC (ThermoFisher Scientific). Chromatography was performed as above but using a 90 min non-linear ACN gradient. Data was collected in Data Dependent Acquisition (DDA) mode using m/z 300–1650 as MS scan range, rCID for MS/MS of the 20 most intense ions. Lockmass of 401.92272 from DMSO was used. Other instrument parameters were: MS scan at 100,000 resolution, maximum injection time 150 ms, AGC target 1E6, CID at 30% energy for a maximum injection time of 150 ms with AGC target of 5000. Dynamic exclusion with of 30 seconds was applied for repeated precursors.

Raw files were analyzed using the MaxQuant platform (61) version 1.6.16.0 searching against the UniProt human database containing 42,434 reviewed, canonical, isoforms entries (June 2019) and a database containing 246 common contaminants. For label-free (LFQ) AE-MS experiments, default search parameters were used with “Label free quantitation” set to “LFQ” and “Match between runs” enabled. In all cases, Trypsin/P cleavage specificity (cleaves after lysine or arginine, even when proline is present) was used with a maximum of 2 missed cleavages. Oxidation of methionine and N-terminal acetylation were specified as variable modifications. Carbamidomethylation of cysteine was set as a fixed modification. A search tolerance of 4.5 ppm was used for MS1 and 20 ppm for MS2 matching. False discovery rates (FDR) were determined through the target-decoy approach set to 1% for both peptides and proteins.

Using the Perseus platform (62) version 1.6.5.0, or 1.6.14.0 or 1.6.15.0, proteins group LFQ intensities were log2 transformed. Values listed as being “Only identified by site,” “Reverse,” or “Contaminants” were removed from the data set. Mitochondrial annotations were imported by matching with the Mitocarta2.0 data set (63) by gene name and/or ENSG identifier. Experimental groups were assigned to each set of triplicates and the number of valid values for row group calculated. For each experiment (containing a control and an enrichment group) rows having less than 3 valid values in the enrichment group were removed and the missing values in the relevant control group imputed to values consistent with the limit of detection. Proteins identified from less than 2 unique peptides were excluded from further analysis. A modified two-sided t test based on permutation-based FDR statistics (62) was performed between the two groups. The negative logarithmic p values were plotted against the differences between the log2 means for the two groups. The significance threshold used for these experiments is noted in the relevant figure legend and supplementary tables. Non-mitochondrial proteins were removed based on their absence from MitoCarta 2.0 (63), before plotting of the data. For SILAC experiments, default search parameters were used with multiplicity set to 2 (Lys8, Arg10) and “Match between runs” enabled. Using the Perseus platform (62) version 1.6.5.0, proteins group normalized H/L ratios were log2 transformed. Label switched samples (L/H) were inverted to KO/Control orientation before this step. Values listed as being “Only identified by site,” “Reverse,” or “Contaminants” were removed from the data set. Mitochondrial annotations were imported by matching with the Mitocarta2.0 data set (63) by gene name and/or ENSG identifier. Experimental groups were assigned to each set of triplicates rows with < 2 valid values for each group removed. Non-mitochondrial proteins were removed based on the absence of a Mitocarta2.0 annotation, and a one sample Student’s two-sided t test was performed within each group. The negative logarithmic p values were plotted against the differences between the mean ratios for each group. A significance threshold (p < 0.05) was used for all experiments. For the topographical heatmap, log2-transformed median SILAC ratios were mapped onto the structure of complex IV (PDB 5Z62).

### Protein Overexpression, Purification, and Characterization

The R39W, Y137C, Y137S, S149I, H119A, M156A and H119A/M156A mutations were introduced into the ORF encoding the COA7 protein in the pGEX-6p-1-COA7 plasmid using a Q5^®^ Site-Directed Mutagenesis kit (New England, BioLabs), according to the manufacturer’s instructions. Forward and reverse primers for all mutant proteins were designed using NEBaseChanger online tool (http://nebasechanger.neb.com/) and are given in **Table S1**. The DNA sequence encoding the variant was amplified *via* PCR followed by ligation and template DNA removal using KLD enzyme mix (containing a kinase, a ligase and DpnI, New England BioLabs^®^).

The pGEX-6p-1 plasmids containing DNA sequence encoding COA7, ^R39W^COA7, ^Y137C^COA7, ^Y137S^COA7, ^S149I^COA7, ^H119A^COA7, ^M156A^COA7and ^H119AM156A^COA7, were individually transformed into *Escherichia coli* strain SHuffle^®^ T7 (New England, BioLabs). Cultures were grown at 30°C in Luria Broth (LB) supplemented with ampicillin (100 μg/mL), chloramphenicol (35 μg/mL) and streptomycin (50 μg/mL) to an OD_600_ of 0.8, induced with isopropyl β-D-1-thiogalactopyranoside (IPTG, 0.2 mM) and harvested after 16 hours (with shaking) at 16°C.

Purified COA7, ^R39W^COA7, ^Y137C^COA7, ^Y137S^COA7, ^S149I^COA7, ^H119A^COA7, ^M156A^COA7 and ^H119AM156A^COA7 proteins were prepared by affinity and size exclusion chromatography. Frozen cell pellets of the GST-fusion proteins were thawed at room temperature and resuspended in PBS (phosphate-buffered saline pH 7.4). Cells were disrupted by passage through a TS series bench top cell disruptor (Constant Systems Ltd) at 35 kpsi. Cell debris were removed by centrifugation (Beckman JLA-25.50, 30000 g, 20 min, 4°C) and the soluble fraction was incubated with glutathione sepharose™ 4B resin (GE Healthcare) equilibrated with PBS. The GST tag was cleaved with PreScission Protease followed by size-exclusion chromatography (SEC; HiLoad 16/600 Superdex 75 pg, GE Healthcare; 20 mM NaHepes pH 7.2, 50 mM NaCl). Cleavage of the N-terminal GST tag introduced five additional residues (GPLGS) to the N-terminus of all proteins. The purified proteins were concentrated to 20 mg/mL before storage at -80°C.

### Sedimentation velocity analysis

COA7 and ^Y137C^COA7 samples were analyzed using an XL-I analytical ultracentrifuge (Beckman Coulter, Fullerton, CA) equipped with an AnTi-60 rotor. Protein samples were loaded in the sample compartment of double-sector epon centrepieces, with buffer (20 mM NaHepes pH 7.5, 50 mM NaCl) in the reference compartment. Radial absorbance data was acquired at 20°C using a rotor speed of 50,000 rpm and a wavelength of 280 nm, with radial increments of 0.003 cm in continuous scanning mode. The sedimenting boundaries were fitted to a model that describes the sedimentation of a distribution of sedimentation coefficients with no assumption of heterogeneity (*c(s)*) using the program SEDFIT (64). Data were fitted using a regularization parameter of p= 0.95, floating frictional ratios, and 100 sedimentation coefficient increments in the range of 0–15 S.

### Differential scanning fluorimetry

The melting temperatures (T_m_) of the COA7, ^R39W^COA7, ^Y137C^COA7, ^Y137S^COA7 and ^S149I^COA7 proteins were measured by differential scanning fluorimetry with purified proteins in 20 mM NaHepes pH 7.2, 50 mM NaCl (1.0 mg/mL). Differential scanning fluorimetry experiments were performed at the CSIRO Collaborative Crystallisation Centre (http://www.csiro.au/C3), Melbourne, Australia.

### Copper binding experiments

Measurements of the copper binding affinities of the COA7 and ATOX1 proteins were performed as previously described (21). Briefly, purified proteins (in 20 mM Tris-Mes pH 8.0) were titrated at various concentrations (1–30 μM) into solutions containing buffer (20 mM Tris-Mes pH 7.0), CuSO_4_ (20 μM), Bcs (200 μM) and NH_2_OH (1 mM). The exchange of Cu(I) from the [Cu^I^Bcs_2_]^3−^ complex to the proteins was monitored by measuring the absorbance of the resulting solutions at 483 nm. The data were analyzed by plotting [Cu^I^Bcs_2_]^3−^ (as determined from the absorbance values at 483 nm) versus protein:copper ratios, and the data fitted using the equation previously described (21, 42).

### Fe^3+^-PPIX and PPIX titrations for the COA7 and ^H119AM156A^COA7 proteins

Fe^3+^-PPIX and PPIX binding assays were performed for the COA7 and ^H119AM156A^COA7 proteins by difference spectroscopy at room temperature and aerobically using a double beam UV-Vis spectrophotometer (Cary 5000 UV-Vis-NIR spectrophotometer) as previously described (43, 44). A baseline was selected by scanning samples from 280 to 750 nm, with buffer (20 mM NaHepes pH 7.2, 50 mM NaCl) in the reference cuvette and 1.0 μM of purified protein (in 20 mM NaHepes pH 7.2, 50 mM NaCl) in the sample cuvette. After the baseline was set, increasing concentrations of Fe^3+^-PPIX (0-10 μM) or PPIX (0-200 μM) were added to both the sample and the reference cuvettes, therefore the readings only reflected the binding of Fe^3+^-PPIX or PPIX to the proteins and not the absorbance of the free ligands. Note that only the titrations with Fe^3+^-PPIX yielded spectra that indicated ligand binding (**Fig. 5 and S10**) and that the PPIX titration was carried out for the COA7 protein only. The data were analyzed by plotting absorbance at 416 nm versus hemin concentration and the data were fitted using equation describing a single binding site model 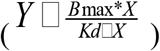 using GraphPad Prism software.

### Fe^3+^-PPIX and PPIX loading of COA7, ^H119A^COA7 and ^M156A^COA7

Purified protein samples were incubated for 1 hour with Fe^3+^-PPIX or PPIX (molar ratio 1:2 protein:ligand) in 20 mM NaHepes pH 7.2, 50 mM NaCl. In order to remove the excess Fe^3+^- PPIX or PPIX from the mixture, the incubated protein sample was applied to a SEC column (10 mL, Bio-Gel P-4 Gel, Bio-Rad). The presence of bound Fe^3+^-PPIX or PPIX in the eluted fractions was detected by UV-Vis spectrophotometry (between 250-700 nm; Cary 300 UV-Vis Absorbance spectrophotometer) and those protein fractions containing bound hemin (with absorption at 416-420 nm) were pooled and concentrated by centrifugal ultrafiltration.

The UV-Vis spectra of Fe^2+^-PPIX-COA7 were recorded (65) following incubation of purified COA7 with one molar equivalent of Fe^3+^-PPIX, and the subsequent addition of 1.0 mM sodium dithionite to reduce all Fe^3+^ to Fe^2+^. UV-Vis spectra of Fe^3+^-PPIX loaded COA7, ^H119A^COA7 and ^M156A^COA7 samples were carried out in presence of sodium cyanide (1.0 mM) (66).

### Metal analysis of the Fe^3+^-PPIX-COA7 complex

The Fe:COA7 stoichiometry of the Fe^3+^-PPIX-COA7 complex was determined by ICP-MS. The Fe^3+^-PPIX-COA7 protein (at various concentrations: 10-100 ppb in buffer: 20 mM NaHepes, pH 7.2, 50 mM NaCl) was digested in 200 μL of 35% HNO_3_ (Merck Suprapur, Australia), diluted to a final volume of 1.0 mL using MilliQ H_2_O and heated at 96°C for 15 min. Insoluble material was removed by centrifugation at 18,000 × g for 25 min. Technical triplicate measurements of samples and controls were analyzed on an Agilent 8900 ICP-MS/MS with a MicroMist nebulizer (Glass Expansion, Australia). Torch positioning, sample depth adjustment and lens optimization were set according to manufacturer recommendations while other instrumental parameters were optimized during a batch-specific user tune. Helium collision gas flow rate of 5 mL/min to minimize polyatomic interferences. Samples were introduced via an integrated automation system (IAS) autosampler (Agilent) using a peristaltic pump. The division of the slope of the standard curve and a plot of Fe concentration versus protein concentration was calculated to yield a value for the Fe:COA7 protein sample.

### Protein Crystallization and Data Collection

Initial high-throughput crystallization experiments using COA7 (20 mg/mL) were performed at the CSIRO Collaborative Crystallisation Centre (http://www.csiro.au/C3), Melbourne, Australia using c3 salty screens (c3_5 and c3_6) by sitting drop vapor diffusion in 96-well plates. Diffraction-quality, plate-like crystals of COA7 grew overnight to a maximum size of 250 x 30 x 20 μ at 20°C by hanging-drop vapor diffusion with drops containing equal volumes (1 μL) of COA7 (20 mg/mL in 20 mM NaHepes pH 7.2, 50 mM NaCl) and crystallization solution (0.23 M sodium Mes pH 6.72, 2.35 M ammonium sulfate, 8.2% (v/v) ethanol and 4.5% (v/v) pentaerythritol ethoxylate (3/4 EO/OH) equilibrated against 500 μL reservoir solution. COA7 crystals were cryoprotected in reservoir solution containing 25% (v/v) glycerol before flash-cooling in liquid nitrogen. Diffraction data were recorded on an EIGER X 16M detector at the Australian Synchrotron on beamline MX2 at a wavelength of 0.954 Å. All data were collected at 100 K and were processed and scaled with XDS (67) and Aimless (68). Data collection statistics are detailed in **Table S3**.

### Structure Solution and Refinement

The crystal structure of COA7 was solved by molecular replacement using the program PHASER (69) from the CCP4 suite (70). Models including residues 33-104 and 110-216 of the HcpC crystal structure (PDB 1OUV) (34) were used as the search models after removal of all water molecules. The structure was refined using REFMAC5 (71), with manual model building and the addition of water molecules carried out in COOT (72). The refinement of the model converged with residuals *R* = 0.20 and *R*_free_ = 0.25 (**Table S3**) and showed excellent geometry as determined by MOLPROBITY (73) (**Table S3**).

## Data Availability

The mass spectrometry proteomics data will be deposited in the ProteomeXchange Consortium via the PRIDE (74) partner repository upon publication. The atomic coordinates and structure factors have been deposited in the Protein Data Bank (https://www.rcsb.org; PDB 7MQZ).

## Acknowledgements

This study was funded by the Australian Research Council (DP140102746 and FT180100397 to MJM), the National Health and Medical Research Council (GNT1165217 to MTR and MJM; GNT1140906 and 1140851 to DAS) and an Australian Government Research Training Program Scholarship to SM. Part of this study was carried out using the MX2 beamline at the Australian Synchrotron, which is part of ANSTO, and made use of the ACRF detector. We thank the beamline staff for their enthusiastic and professional support. We thank Dr. Yee-Foong Mok at the Melbourne Protein characterization facility, The Bio21 Molecular Science and Biotechnology Institute, The University of Melbourne for performing the AUC experiments and data analyzes. We thank the Bio21 Mass Spectrometry and Proteomics Facility for the provision of instrumentation, training, and technical support. We thank Daniel Machell for Pymol and coding assistance and Dr Katie Ganio and Prof. Christopher McDevitt for the ICP-MS analysis. Finally, we thank Mrs Mahnaz Dideh Var for her extraordinary support with the structural biology aspects of this study.

