## Supplementary Information for "Mitochondrial COA7 is a heme-binding protein involved in the early stages of complex IV assembly"

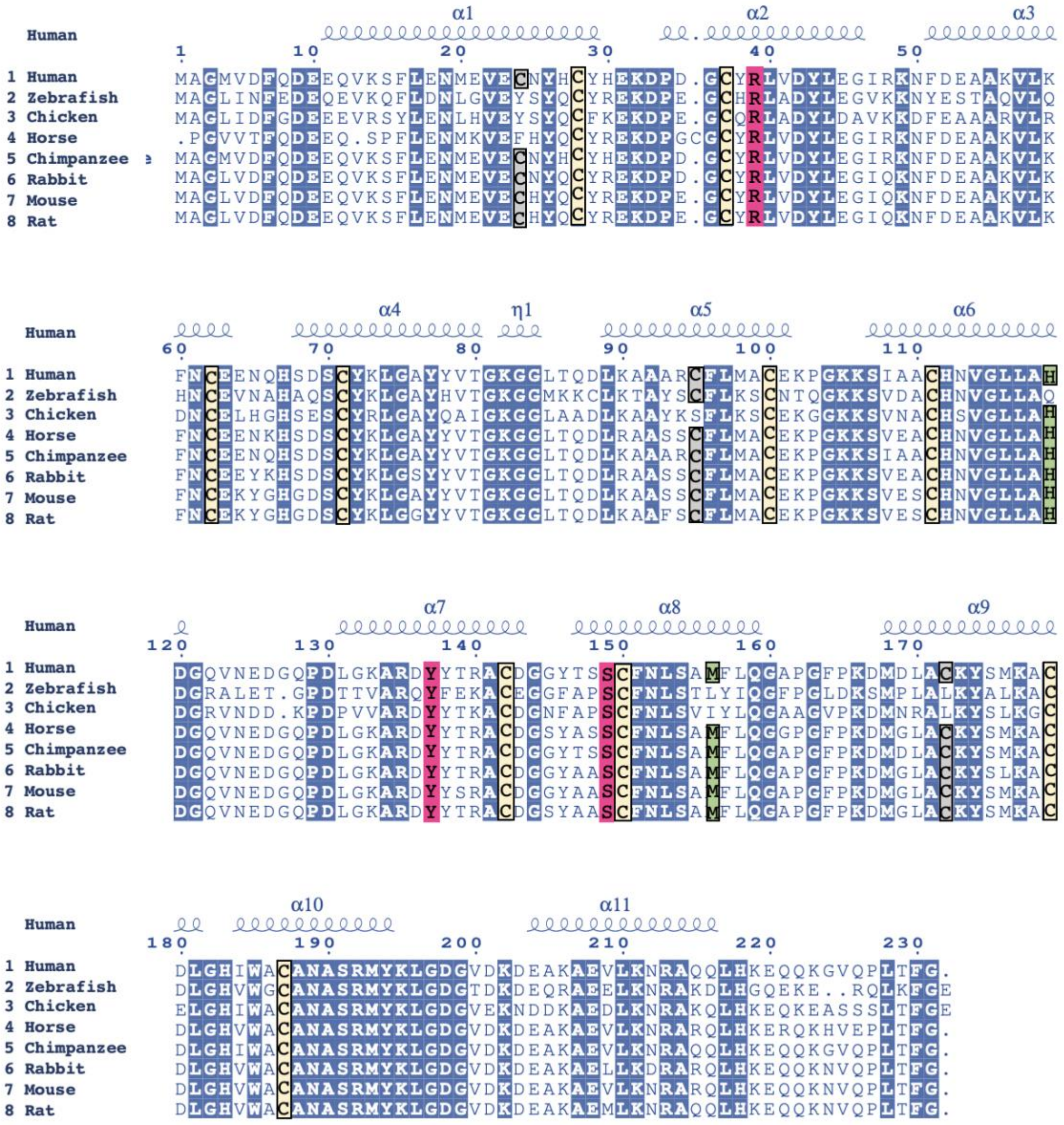

**Figure S1.** Sequence alignment between COA7 proteins across metazoans using *Clustal omega* and rendered with *ESPript* (1, 2). Numbering is relative to human COA7. Conserved cysteine residues are highlighted in yellow. Pathogenic mutations are highlighted in pink. Gray boxes indicate cysteine residues that are not conserved across eukaryotes. Residues His119 and Met156 (human numbering) are highlighted in green. All remaining conserved residues are highlighted in blue.

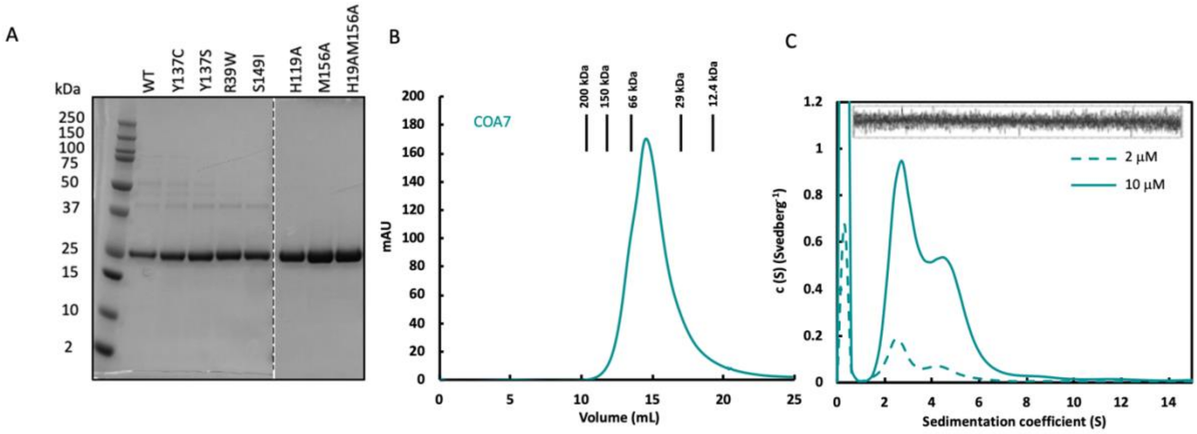


**Figure S2.** Purification and characterization of the COA7 proteins. (A) Purified COA7, $^{Y137C}$COA7, $^{Y137S}$COA7, $^{R39W}$COA7, $^{S149I}$COA7, $^{H119A}$COA7, $^{M156A}$COA7, and $^{H119AM156A}$COA7 proteins were analyzed by SDS-PAGE. (B) SEC elution profile of purified COA7 relative to indicated protein standards. (C) Analytical ultracentrifugation sedimentation velocity analysis of the purified COA7. Sedimentation velocity continuous size [*c(s)*] distribution best fit for COA7 measured at protein concentrations of 2 μM (dashed lines, teal) and 10 μM (solid lines, teal).

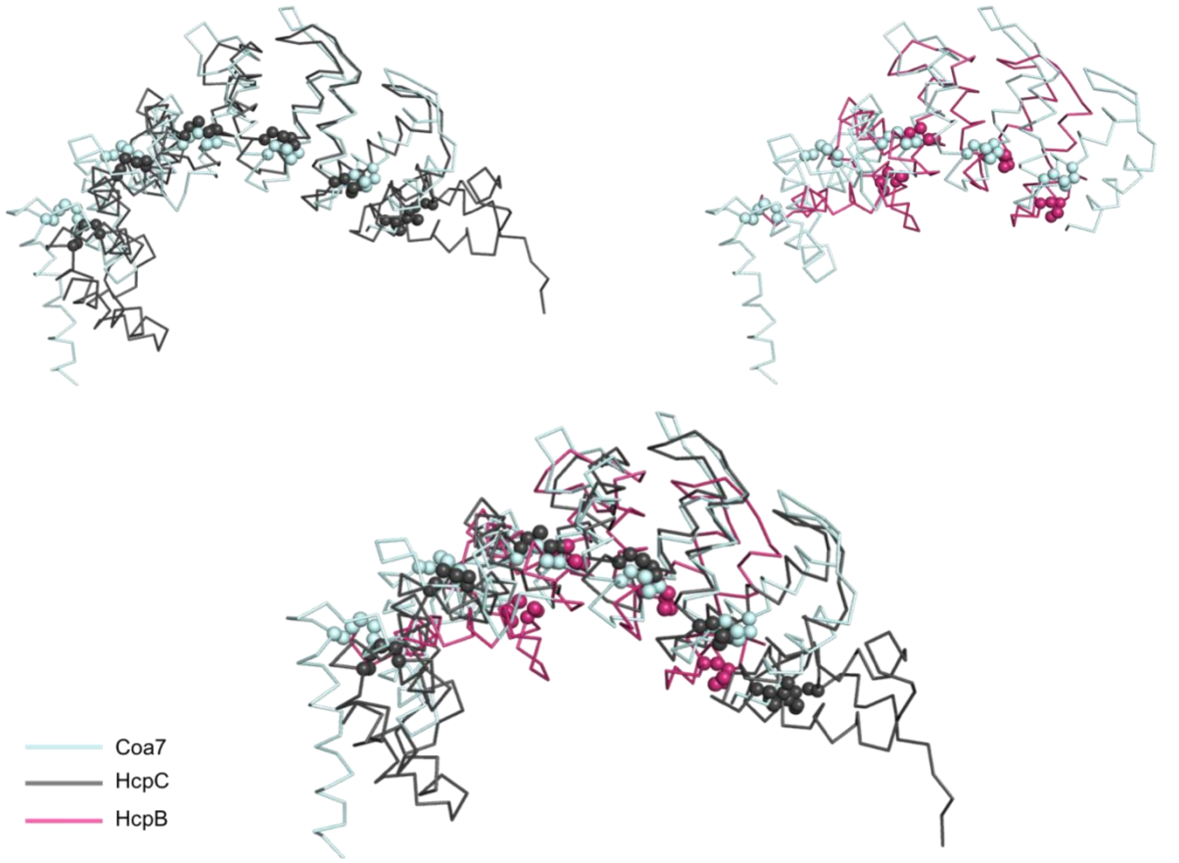


**Figure S3.** Comparison of the COA7 (cyan), HcpB (pink; PDB 1KLX) (3) and HcpC (gray; PDB 1OUV) (4) structures. The overall fold of COA7 is similar to Hcp proteins composing of disulphide-bridged $\alpha/\alpha$ repeats. Cysteine residues involved in intramolecular disulfide bonds formation are shown as spheres.

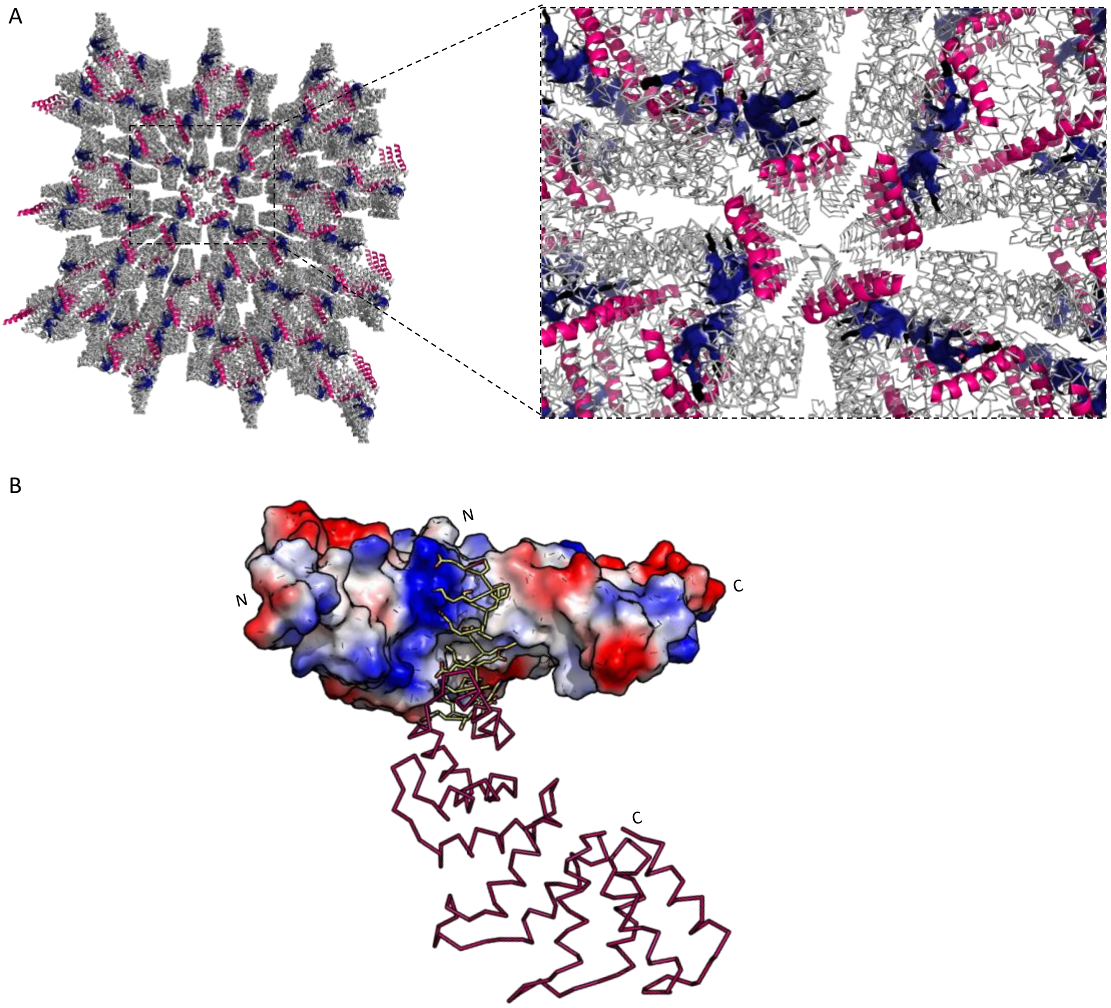


**Figure S4.** COA7 crystal packing. (*A*) The application of crystallographic symmetry operators reveals that in the crystal, the cleft in the concave face of the molecule is occupied by the N-terminal helix (pink) of a neighboring molecule of COA7, such that the crystal is composed of an infinite network of protein-protein interactions (*B*) Interactions observed in COA7 crystal contact. The molecular surface of COA7 is colored according to the electrostatic potential (negative potential, red; positive potential, blue). The main chain of the symmetry-related COA7 molecule is shown as a ribbon. Side-chains (yellow sticks) of the N-terminal helix of the symmetry-related COA7 molecule interact with the concave surface through hydrophobic and electrostatic interactions.

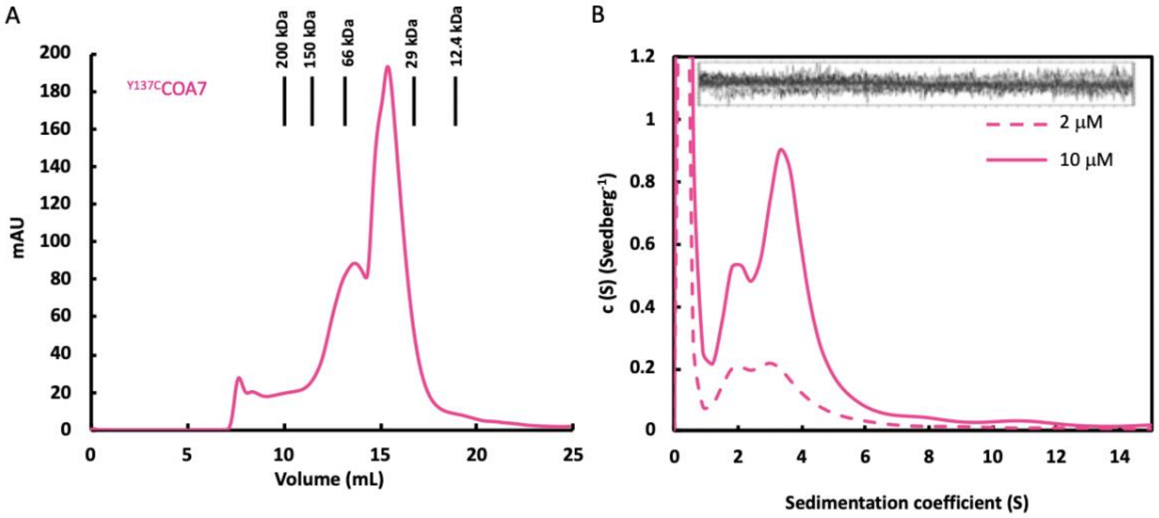


**Figure S5.** Purification and characterization of the $^{Y137C}$COA7 protein. (*A*) Elution profile of purified $^{Y137C}$COA7 relative to indicated protein standards analyzed by size-exclusion chromatography. (*B*) Analytical ultracentrifugation sedimentation velocity analysis of the purified $^{Y137C}$COA7. Sedimentation velocity continuous size [*c(s)*] distribution best fits for $^{Y137C}$COA7 measured at protein concentrations of 2 µM (dashed lines, pink) and 10 µM (solid lines, pink).

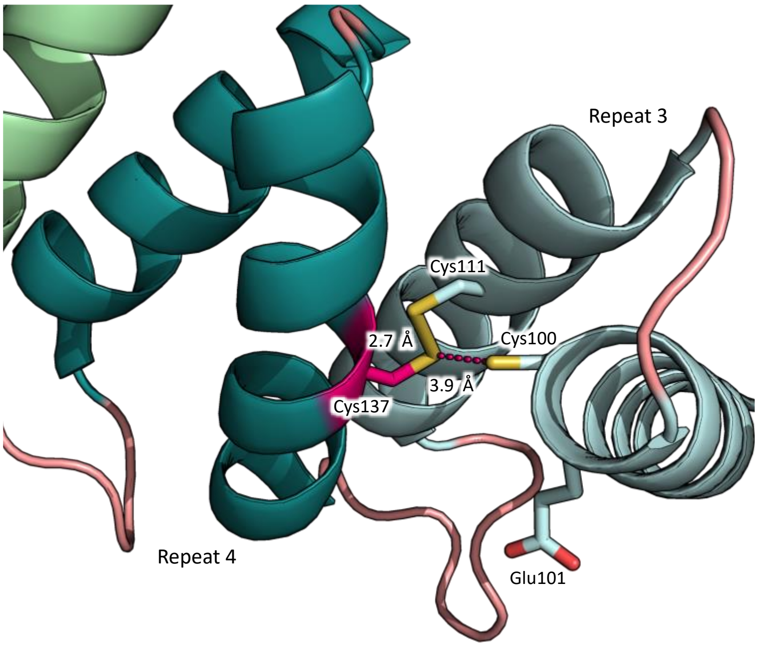


**Figure S6.** The Y137C pathogenic mutation may lead to the disruption of the Cys100-Cys111 disulfide bond. The COA7 structure shows that the formation of an alternative Cys137-Cys111 disulfide bond would be possible in the presence of the Y137C mutation (predicted Cys137 SG-Cys111 SG distance = 2.7 Å). However, the predicted distance between the thiolate groups of introduced Cys137 and residue Cys100 is 3.9 Å, indicating that a Cys137-Cys100 bond would not form in the absence of a major structural reorganisation.

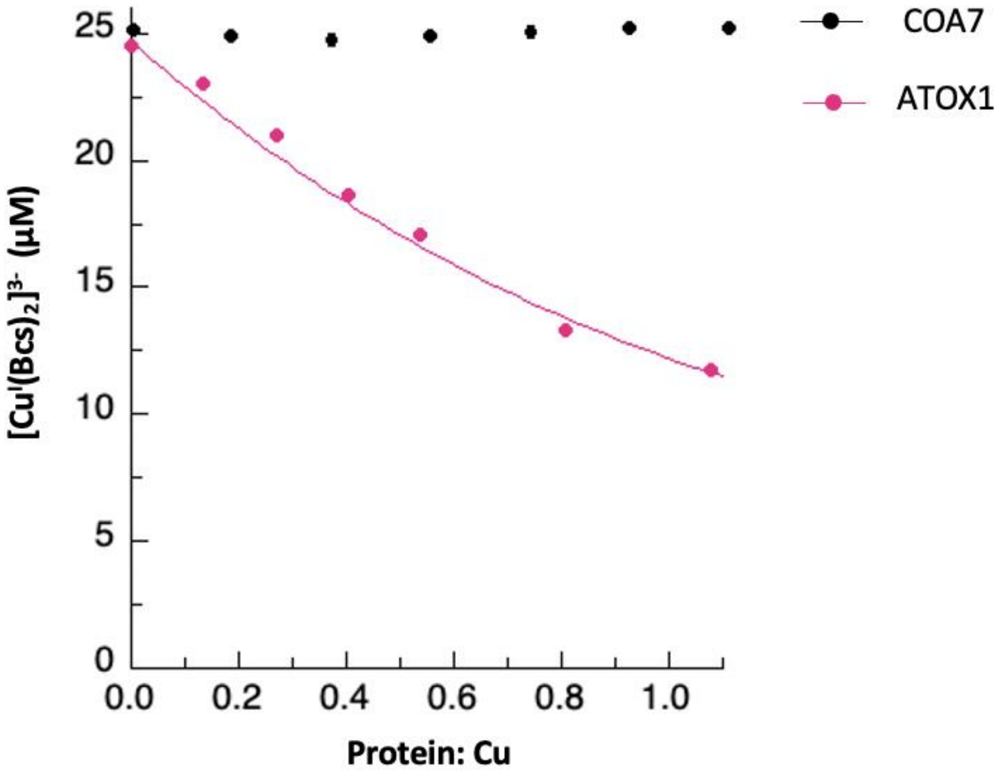


**Figure S7.** COA7 exhibits no copper binding. Known concentrations (1–30 μM) of recombinant COA7 and ATOX1 were added to a complex of $[Cu^IBcs_2]^{3-}$ (24 μM), and the absorbance of the solution was measured at 483 nm. The data were analyzed using plots of $[Cu^IBcs_2]^{3-}$ versus COA7/ATOX1:copper ratio, and the data fit as previously described (5, 6).

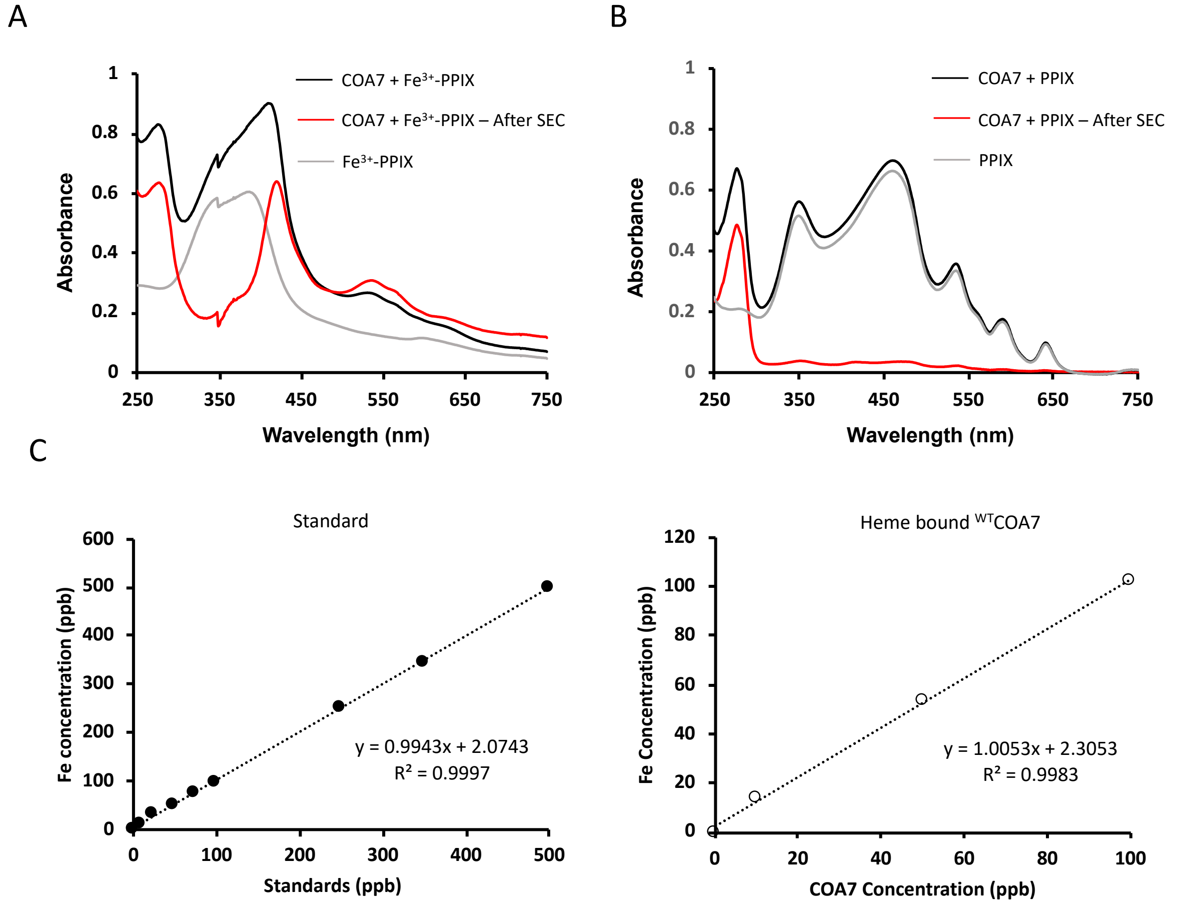


**Figure S8.** COA7 co-elutes from SEC with $Fe^{3+}$-PPIX but not with PPIX. *(A)* Free hemin has a UV-Vis absorption spectrum with maxima at 365 and 385 nm (gray), whereas bound hemin has a UV-Vis absorption spectrum with maxima at 412 and 420 nm. The UV-Visible spectrophotometry data show that before SEC sample contains a broad peak (black); corresponding to both the presence of free hemin (385 nm) and bound hemin with a Soret maximum at 416 nm. However, after SEC the spectrum of the resultant protein shows a Soret band with maxima at 416 nm only indicating the presence of the heme-COA7 complex. *(B)* Comparison the spectra of PPIX in the absence and presence of COA7 (gray and black, respectively) shows limited differences. The sample following SEC (red) shows minimal features in the visible region of the spectrum, indicating minimal/no PPIX-protein binding. *(C)* Determination of the Fe content of heme bound COA7 by ICP-MS. A standard curve (left) was used to determine the Fe concentration of the heme-COA7 (right).

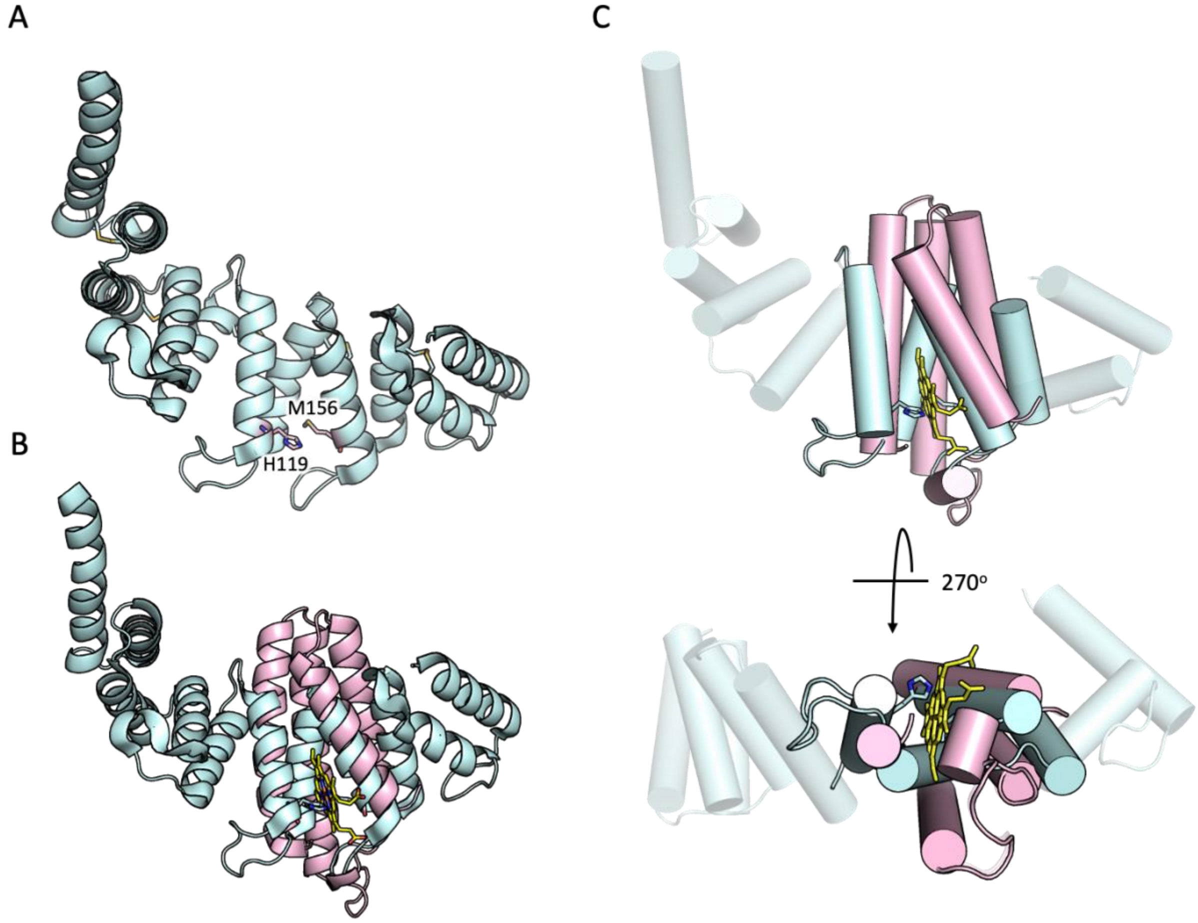


**Figure S9.** Examination of the structure of COA7 reveals a possible heme binding site located between repeats 3 and 4, with residues His119 and Met156 in comparable positions to those that act as iron ligands in the cytochrome $b_{562}$ from *E. coli*. *(A)* Histidine 119 and Methionine 156 are shown as pink sticks on the COA7 structure. *(B)* Superposition of the crystal structure of COA7 (cyan) with that of cyt $b_{562}$ (pink; PDB 256B) (7). The heme cofactor in cyt $b_{562}$ is indicated with yellow sticks. *(C)* The same superposition as in (B) with α-helices represented as cylinders.

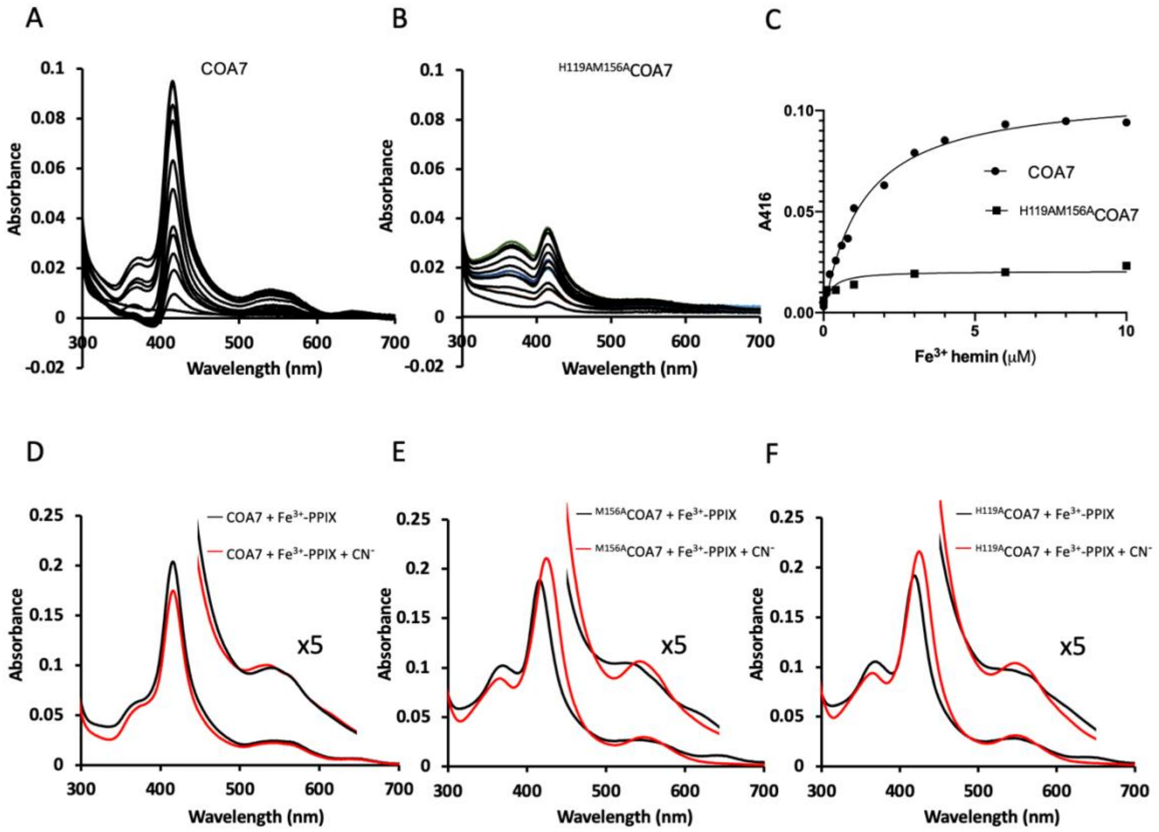


**Figure S10.** *(A)* Heme binding to COA7. *(B)* Heme binding to $^{H119AM156A}$COA7. Difference absorption spectra of $^{H119AM156A}$COA7 (1 μM) with increasing concentrations of $Fe^{3+}$ hemin (from 0 to 10 μM). *(C)* The binding curve was fitted to the data from *A* and *B* using an equation describing a single binding site (Y = Bmax * X/KD + X) with GraphPad Prism. Unlike COA7, no significant binding of $Fe^{3+}$-PPIX to $^{H119AM156A}$COA7 can be detected. *(D)* UV-Vis spectra of $Fe^{3+}$-PPIX loaded COA7 in the absence (black) and presence (red) of 1 mM sodium cyanide ($CN^-$). *(E)* UV-vis absorption spectra of $Fe^{3+}$-PPIX loaded $^{M156A}$COA7 in the absence (black) and presence (red) of 1 mM sodium cyanide ($CN^-$). *(F)* UV-vis absorption spectra of $Fe^{3+}$-PPIX loaded $^{H119A}$COA7 in the absence (black) and presence (red) of 1.0 mM sodium cyanide ($CN^-$).

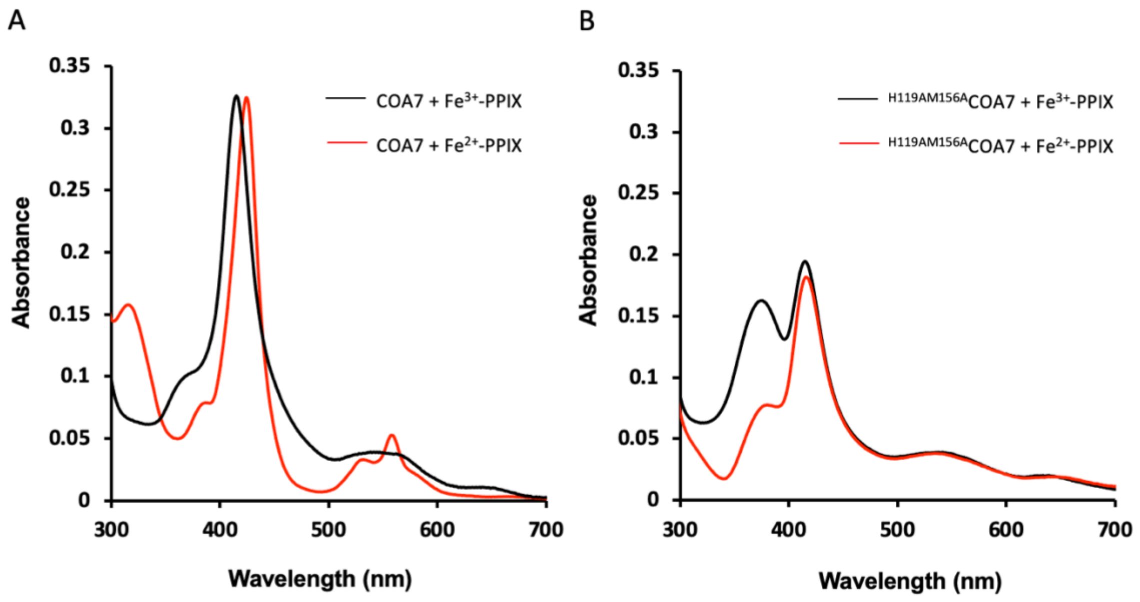


**Figure S11.** Comparison the spectra of COA7 and $^{H119AM156A}$COA7 incubated with one equivalent of $Fe^{3+}$-PPIX (black) and following reduction with sodium dithionite ($Fe^{2+}$-PPIX red). The absence of alpha and beta bands at ~550 nm (*B*, red) and the weak Soret peak for the $^{H119AM156A}$COA7 (~420 nm) indicates a lack of axial coordination to the iron by amino acid ligands.

**Table S1. Forward and reverse primers for all COA7 mutant proteins.**

| Protein | Forward primer | Reverse primer |
|---|---|---|
| R39W | TGGTTGCTACTGGCTGGTGGACT | TCCGGGTCCTTCTCGTGATAG |
| H119A | TCTGCTGGCGGCGGATGGTCAGGTTAACGAGGACG | CCCACGTTGTGGCACGCC |
| M156A | CCTGAGCGCGGCGTTTCTGCAGG | TTGAAGCAGCTGCTGGTATAG |
| Y137C | GGCGCGTGACTGCTATACCCGTG | TTGCCCAGATCCGGTTGAC |
| Y137S | GGCGCGTGACAGCTATACCCGTG | TTGCCCAGATCCGGTTGAC |
| S149I | CTATACCAGCATCTGCTTCAACCTGAGC | CCACCATCGCACGCACGG |

**Table S2. Hydrodynamic properties of the COA7 and $^{Y137C}$COA7 proteins.**

| Protein | $M_r$ (Da)[a] | s (S)[b] | M (kDa)[c] | $f/f_0$ |
|---|---|---|---|---|
| COA7 | 25649 | 1.9 | 26 | 1.3 |
| | | 3.5 | 54 | |
| $^{Y137C}$COA7 | 25620 | 2.1 | 28 | 1.2 |
| | | 3.5 | 64 | |

[a] Molecular mass of the proteins were determined from the protein primary sequences.

[b] Modal sedimentation coefficients were taken from the ordinate maximum of the *c(s)* distribution best fit for sedimentation velocity data generated at initial protein concentrations of 10 µM for COA7, $^{Y137C}$COA7 proteins (Fig. 4E and F, respectively).

[c] Molar masses were determined from the *c(M)* distribution best fit (data not shown).

**Table S3. Data collection and refinement statistics**

| **Data collection** | |
|---|---|
| Crystal | COA7 |
| Wavelength (Å) | 0.953654 |
| Temperature (K) | 100 |
| Diffraction source | Australian Synchrotron (MX2) |
| Detector | EIGER X 16M |
| Space group | $I4_1$ |
| *a*, *b*, *c* (Å) | 100.01, 100.01, 50.34 |
| α, β, γ (°) | 90.00, 90.00, 90.00 |
| Resolution range (Å) | 44.97- 2.39 (2.48-2.39) |
| Total No. of reflections | 68093 |
| No. of unique reflections | 10025 |
| Completeness (%) | 99.7 (96.9) |
| Redundancy | 6.8 (6.7) |
| $CC_{1/2}$ | 0.99 (0.81) |
| $\langle I/\sigma(I)\rangle$ | 10.5 (1.6) |
| *R*merge (%) | 9.3 (73.0) |
| *R*pim (%) | 3.9 (30.1) |
| **Refinement statistics** | |
| Resolution range (Å) | 70.72-2.39 (2.448-2.39) |
| No. of reflections, working set | 949720 |
| No. of reflections, test set | 521 |
| $R_{work}$ (%) | 20.52 (29.1) |
| $R_{free}$ (%) | 25.47 (35.9) |
| Rmsd bond lengths (Å) | 0.010 |
| Rmsd bond angles (°) | 1.301 |
| Ramachandron[†] | |
| Favored, % | 99.1 |
| Allowed, % | 100 |
| PDB ID code | 7MQZ |

*Values in parenthesis are for highest-resolution shell.

[†]Calculated using MolProbity (8).
